# Lethal COVID-19 Associates With RAAS-Induced Inflammation For Multiple Organ Damage Including Mediastinal Lymph Nodes

**DOI:** 10.1101/2023.10.08.561395

**Authors:** Joseph W. Guarnieri, Michael J. Topper, Katherine Beigel, Jeffrey A. Haltom, Amy Chadburn, Justin Frere, Julia An, Henry Cope, Alain Borczuk, Saloni Sinha, Christine Lim, JangKeun Kim, Jiwoon Park, Cem Meydan, Jonathan Foox, Christopher Mozsary, Yaron Bram, Stephanie Richard, Nusrat J. Epsi, Brian Agan, Josh Chenoweth, Mark Simons, David Tribble, Timothy Burgess, Clifton Dalgard, Mark T. Heise, Nathaniel J. Moorman, Victoria K. Baxter, Emily A. Madden, Sharon A. Taft-Benz, Elizabeth J. Anderson, Wes A. Sanders, Rebekah J. Dickmander, Gabrielle A Widjaja, Kevin Janssen, Timmy Lie, Deborah Murdock, Alessia Angelin, Yentli Soto Albrecht, Arnold Olali, Zimu Cen, Joseph Dybas, Waldemar Priebe, Mark R. Emmett, Sonja M. Best, Maya Kelsey Johnson, Nidia S. Trovao, Kevin B. Clark, Victoria Zaksas, Rob Meller, Peter Grabham, Jonathan C. Schisler, Pedro M. Moraes-Vieira, Simon Pollett, Christopher E. Mason, Eve Syrkin Wurtele, Deanne Taylor, Robert E. Schwartz, Afshin Beheshti, Douglas C. Wallace, Stephen B. Baylin

## Abstract

Lethal COVID-19 outcomes are most often attributed to classic cytokine storm and attendant excessive immune signaling. We re-visit this question using RNA sequencing in nasopharyngeal and 40 autopsy samples from COVID-19-positive and negative individuals. In nasal swabs, the top 100 genes which significantly correlated with COVID-19 viral load, include many canonical innate immune genes. However, 22 much less studied "non-canonical" genes are found and despite the absence of viral transcripts, subsets of these are upregulated in heart, lung, kidney, and liver, but not mediastinal lymph nodes. An important regulatory potential emerges for the non-canonical genes for over-activating the renin-angiotensin-activation-system (RAAS) pathway, resembling this phenomenon in hereditary angioedema (HAE) and its overlapping multiple features with lethal COVID-19 infections. Specifically, RAAS overactivation links increased fibrin deposition, leaky vessels, thrombotic tendency, and initiating the PANoptosis death pathway, as suggested in heart, lung, and especially mediastinal lymph nodes, with a tightly associated mitochondrial dysfunction linked to immune responses. For mediastinal lymph nodes, immunohistochemistry studies validate the transcriptomic findings showing abnormal architecture, excess fibrin and collagen deposition, and pathogenic fibroblasts. Further, our findings overlap findings in SARS-CoV-2 infected hamsters, C57BL/6 and BALB/c mouse models, and importantly peripheral blood mononuclear cell (PBMC) and whole blood samples from COVID-19 patients infected with early variants and later SARS-CoV-2 strains. We thus present cytokine storm in lethal COVID-19 disease as an interplay between upstream immune gene signaling producing downstream RAAS overactivation with resultant severe organ damage, especially compromising mediastinal lymph node function.

## INTRODUCTION

Coronavirus Disease 2019 (COVID-19), caused by the single-stranded RNA virus, Severe Acute Respiratory Syndrome Coronavirus 2 (SARS-CoV-2), is responsible for extensive global morbidity and mortality^1^. Various studies have probed the molecular underpinnings of patient responses to the virus for the most severe consequences. These studies employed RNAseq analyses of nasal swabs and blood cells but less often visceral tissues^2,3^. Initial RNAseq analysis of nasal swabs and cell models showed that a deficit in responses of classic interferon (IFN) driven viral defense genes might account for overactive innate immune responses in infected patients. However, subsequent analyses indicated IFN genes are robustly upregulated, especially in the nasopharynx^2,4^. RNAseq and metabolomic analysis of bronchial alveolar fluid (BALF), whole blood, and serum collected from severe COVID-19 patients displayed excessive cytokine levels correlating with COVID-19 severity^3,5,6^. However, no studies have investigated the effect of COVID-19 on the immune system in a full spectrum of autopsy organs, which is imperative to understanding how deficits in innate and adaptive immune responses can result in tissue damage leading to lethal outcomes^7^.

Many critically ill patients with COVID-19 and Post-acute Sequelae of COVID-19 (PASC) have shown damage to multiple organ systems, including severe lung, kidney, and cardiac injury, liver dysfuction^8^, and lymphedema^9^. One possible explanation for this widespread organ damage might be a chronic overactivation of RAAS^10,11^. RAAS normally regulates blood pressure but, when chronically overactivated, triggers a cascade of inflammatory tissue-damaging events which are prominent in HAE, an organ dysfunction disorder with similarity to severe COVID-19^10^. Proper regulation of RAAS activity is controlled by ACE2, the receptor for cellular entry of SARS-CoV-2, and a key upstream event for RAAS overactivation is the reduction of ACE2^10^. Indeed, one BALF RNAseq study of severe COVID-19 patients reported altered levels of two RAAS components, bradykinin and hyaluronic acid, and decreased levels of SERPING1, which normally inhibits complement activity. The investigators suggested severe COVID-19 reflects a disease driven by "bradykinin storm," wherein a vascular leak syndrome results in severe pulmonary inflammation with respiratory compromise and risk of thrombosis^12^.

Previous studies have reported mitochondrial dysfunction in COVID-19 revealing that mitochondrial defects activate the innate immune system through mitochondrial oxidative phosphorylation (OXPHOS) inhibition inducing elevated mitochondrial reactive oxygen species (mROS)^7^. The result is release of mitochondrial DNA (mtDNA) and mitochondrial double-stranded RNA (dsRNA) into the cytosol^13,14^. Cytosolic mt-DNA and -dsRNA can be detected by innate immune sensors, resulting in the activation of an innate immune response and initiation of cell death pathways^6,13–17^. Hence, viral inhibition of mitochondrial bioenergetics may play an important role in the activation and intensity of SARS-CoV-2-induced inflammation.

We now rigorously probe how damage to multiple organs may occur in patients with lethal outcomes from COVID-19 by comparing RNAseq in nasopharyngeal samples and autopsy tissues and performing histopathology measurements to validate immune defects correlated with RAAS activation as suggested by our transcriptomic findings. We link transcriptional changes to the critical role of mitochondrial damage which induces this organelle to act as a gateway to activation of the innate immune response and integrated stress response (ISR). Further, we validate our findings in murine models of SARS-CoV-2 infection and through examination of peripheral blood mononuclear cells (PBMCs) and whole blood samples from patients infected with both early and later SARS-CoV-2 variants.

A major finding is that in nasopharyngeal samples, the top-upregulated innate immune genes and adaptive immune genes are linked as upstream signals to extensive gene alterations corresponding to RAAS-overactivation. Our results reveal a complex interplay between upstream immune gene signaling and downstream RAAS activation to induce lethal damage to multiple organ systems and an expanded view of what cytokine storm constitutes. While others have suggested this RAAS paradigm^11^, detailed primary data for the full spectrum of events has been lacking in the COVID-19 literature. Our present work now suggests involvement of many genes not previously focused upon, highlighting components of overactive RAAS signaling such as PANoptosis, a process mediating multiple modes of cell death. A seminal finding, which we document with laboratory studies, is how the RAAS component of excess fibrin deposition and associated collagen deposition is associated with fibroblast infiltration of the mediastinal lymph nodes, correlating with altered lymph node architecture, histological indicators of fibrosis, and immune cell abnormalities, which may all contribute to a reduced immune response integral to COVID-19 patient mortality.

## RESULTS

### Sample collection for RNAseq analysis and SARS-CoV-2 viral load

We quantified the relative expression level of human host and SARS-CoV-2 transcripts from RNAseq data collected from 735 nasopharyngeal and in 40 lung, heart, kidney, liver, and mediastinal lymph node autopsy samples SARS-CoV-2 positive and negative individuals. Our analysis revealed high levels of SARS-CoV-2 transcripts in the human nasopharyngeal samples with significantly lower detectable SARS-CoV-2 transcripts in the human autopsy samples (**Table S1**).

As we could not correlate analyses with precise time points after COVID-19 diagnosis, we analyzed expression levels of host genes and SARS-CoV-2 genes in RNAseq data from SARS-CoV-2 infected Syrian hamsters and C57BL/6 and BALB/c mouse models, where sample collection timing is controlled. In the hamster, heart, kidney, brain, and lung, samples were collected when viral titers peaked in the lung. In the mice, lung samples were collected after peak viral titers (**Table S1**). SARS-CoV-2 gene transcripts were highest in the hamster lungs, followed by the BALB/c and C57BL/6 mouse lungs; while significantly lower levels of SARS-CoV-2 transcripts were detected in the hamster heart, kidney, or brain (striatum, cerebellum, olfactory bulb). Together, these analyses enabled us to assess COVID-19 progression early in infection in the nasopharyngeal and hamster samples, mid-infection in the mice, and after the virus had been cleared in the human autopsy tissues.

### Early-Stage Transcript Changes in Human Nasopharyngeal Samples

#### Individual responses to SARS-CoV-2 infection

To survey individual gene expression responses to SARS-CoV-2, a z-score vector of gene expression for positive samples versus PCR “None” group was generated per nasopharyngeal sample, and the ranked z-score vector was used for gene enrichment analysis versus selected gene sets. **Figure 1A** shows the NES results within the Hallmark IFNα gene set and reveals that for most immune-related gene sets, the group effects were significantly linear for the amount of PCR-detected SARS-CoV-2 between groups.

**Fig 1.**
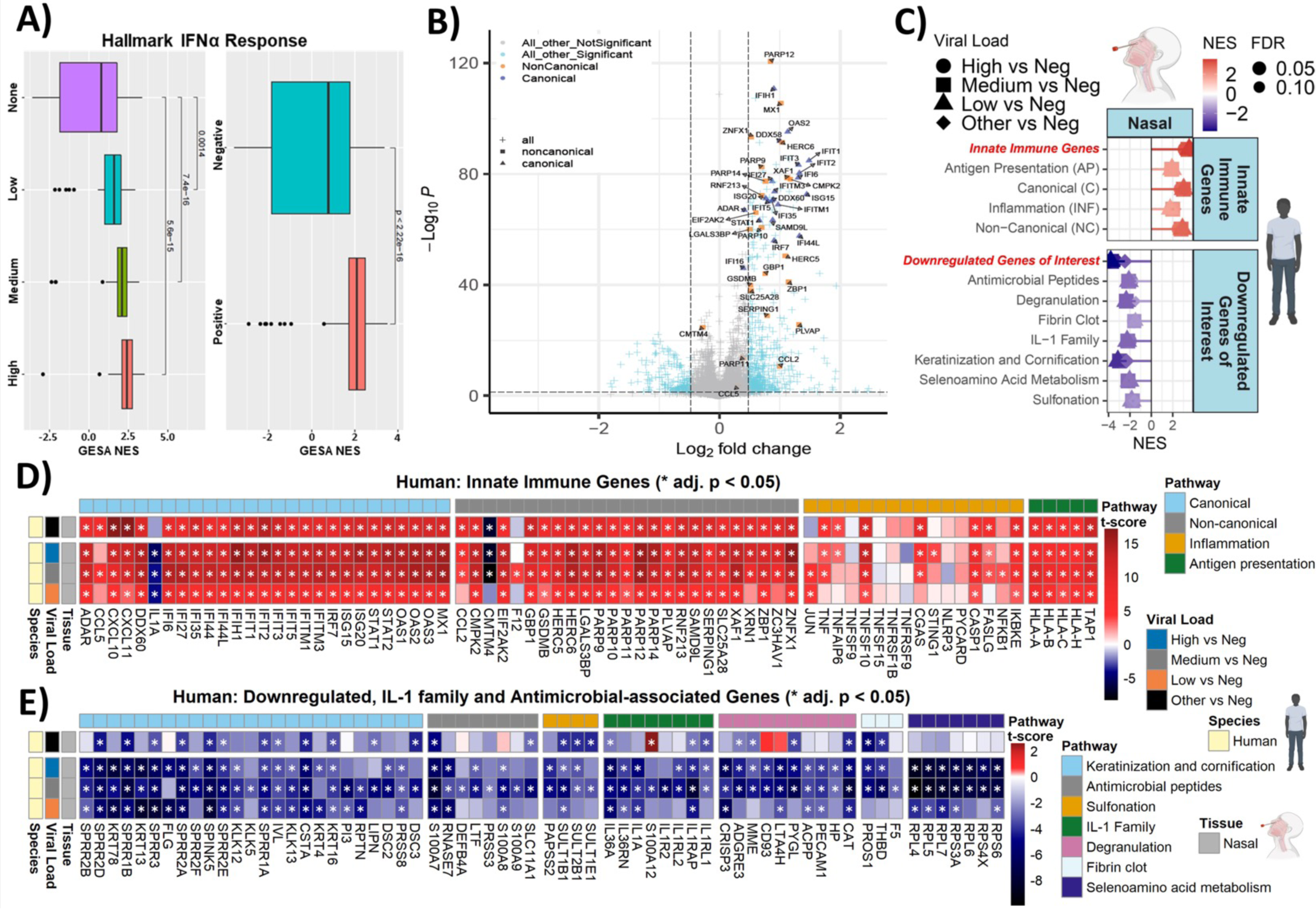
Early-strage innate immune gene expression in nasopharyngeal samples from COVID-19 patients. **A)** Box and whisker plot of hallmark interferon alpha response gene sets determined by fGSEA for nasopharyngeal samples ranked by NES (nominal enrichment score). **B)** Volcano plots of nasopharyngeal samples with high, SARS-C0V-2 RNA copy levels at collection. **D-E)** Heatmaps of t-score statistics when comparing viral load versus negative patient samples for specific **D)** innate immune genes and **E)** downregulated genes of interest.

#### Upregulated, canonical innate immune genes

We next determined relative gene expression levels from RNAseq data in SARS-CoV-2 positive and negative nasopharyngeal samples (**Fig. 1B-E**) fitted for viral load using curated gene lists to characterize the highest up- and downregulated genes in samples from COVID-19 infected versus negative patients to guide subsequent analyses.

Twenty-nine of the most differentially upregulated genes, which we term "canonical" innate immune genes; that are the well known IFN-driven and sensitive (ISG) genes for proteins processing viral dsRNA including *IFIH1, IFIT2, OAS1-3, STAT2, ISG20, DDX60*, and *ISG15* among others (**Figs. 1B-D**). These canonical genes again stratify with increasing viral load (**Figs. 1D**) and with induction of the IFN alpha GESA pathway (**Fig. 1A**). All canonical genes were upregulated except for *IL1A,* which was downregulated (**Fig. 1D**). Importantly, IL1A is released from infected cells to signal immune clearance^18^; thus, its downregulation could prevent these cells from being cleared. Additionally, several genes for protein assembly factors for surface expression of the histocompatibility complex (MHC) 1 allowing complex antigen recognition, and NF-κB and TNF-related genes were also upregulated.

#### Non-canonical innate immune genes

Among the most differentially upregulated genes are 22 ISGs not as generally, or even noted in COVID-19 papers but clearly designated in literature as IFN, ISG genes. We termed these "non-canonical" innate immune genes (**Figs. 1B-D**). These genes also stratified with increasing viral load and were all significantly upregulated except *CMTM4*, whose downregulation could promote T-cell exhaustion^19^, which is common in severe COVID-19^20^. Importantly, these *genes* play important roles throughout our narrative and some key aspects include: **1**) *SERPING1* is crucial for controlling complement pathway activation, a key event in severe tissue-damaging inflammation^10,21^; **2**) *PLVAP*, an endothelial cell-specific membrane protein required for controlling microvascular permeability and barrier function of gut endothelium^22^; when deleted in mice triggers fluid and protein leakage into surrounding tissues and premature death^23^. **3**) *RNF213* encodes a C3HC4-type RING-finger domain that facilitates Class I MHC antigen processing and presentation and Wnt-signaling inhibition^24,25^. **4**) *HERC5/6* both act as anti-viral IFN genes that intersect with ISG15, an ISGylation ligase-type enzyme that targets viral-RNA/proteins for degradation^26–28^. **5**) Five poly (ADP-Ribose) polymerase (PARP) genes are upregulated (**Fig. 1D**), which **r**espond to cellular stressors, including oxidative stress, DNA damage, and pathogen infection^29^. Most PARP genes are ribosylases and MARylases, blocking the translation of viral RNAs and targeting viral proteins for degradation^29^. Notably, several upregulated non-canonical genes, including *ZNFX1, ZBP1, GSDMB, XAF1,* and *CMPK2,* utilize mitochondria to activate the innate immune system. Among these the top one, *ZNFX1,* encodes a zinc finger helicase, which upon viral infection recognizes viral dsRNA and shuttles to the outer mitochondrial (Mt) membrane to restrict viral replication ^30,31^. *ZBP1* has the potential to induce multiple modes of cell death, including apoptosis, pyroptosis, necroptosis, and panoptosis^32–35^ (**Fig. 1D**). *GSDMB* is an integral member of the gasdermin (GDSM) family of death mediators^36^; *XAF1* mediates TNFα-induced apoptosis^37^; *CMPK2* stimulates Mt DNA synthesis, a key dynamic for inducing inflammasome signaling^13,38^

#### Downregulated IL-1 family and antimicrobial response-associated genes

Eighty-three down-regulated genes in the nasopharyngeal samples (**Figs. 1B, E**) encompass seven functional categories including **1**) four sulfonation-associated genes (SULT2B1, SULT1E1, SULT1B1, PAPSS2); **2**) seven selenoamino acid metabolism (RPL4/5/6/7, RPS3A/4X/S6) genes; **3**) sixteen cornification and keratinization-associated genes; **4**) eight antimicrobial peptide/degranulation genes; and four IL-1 family genes. The latter two categories have been previously suggested to be cytokine storm components during SARS-CoV-2 infection^39^ and sulfonation and selenoamino acid metabolism gene upregulation indicates the potential involvement of keratinocytes in SARS-CoV-2 nasopharyngeal pathogenesis.

### Later-Stage Transcript Changes in Human Autopsy Tissues and Rodent Antecedents

To better understand inflammation processes that underlie severe COVID-19, we compared transcription in the nasopharyngeal samples to a multitude of autopsy tissues and further validated key findings in SARS-CoV-2 infected-hamster and -C57BL/6 and -BALB/c mice.

#### Comparison of canonical, inflammation, and antigen presentation genes

Despite the absence of SARS-CoV-2 transcripts, activation of immune and inflammatory genes is seen in all assessed tissues except for liver and lymph nodes. In total, twenty-nine canonical immune genes highly expressed in nasopharyngeal samples retain significant upregulation in the heart (17), kidney (17), and lung (14) (**Figs. 2A, B**). In sharp contrast, in both liver and lymph nodes, only 2 genes are upregulated and 7 are downregulated including those encoding Type 1 (HLA) associated genes (**Fig. 2A**). In lung, heart, and kidney as compared to a lesser extent in the nasopharyngeal samples, there is robust upregulation of death-associated TNF-superfamily (*TNFSFs*) genes that accompany IFN-signaling^40^ (**Fig. 2A**) including *TNSF10* in heart, and *TNSF1/5/9* and *TNFAIP6* in the lung. Additionally, *CXCL10/11* are upregulated in lung, heart, kidney, and nasopharyngeal samples (**Fig. 2A**). CXCL10 is a potent inflammatory cell chemotactic chemokine known to be increased in the serum of severe COVID-19 patients^41^ and mouse lungs infected with SARS-CoV-2^42^.

**Fig 2.**
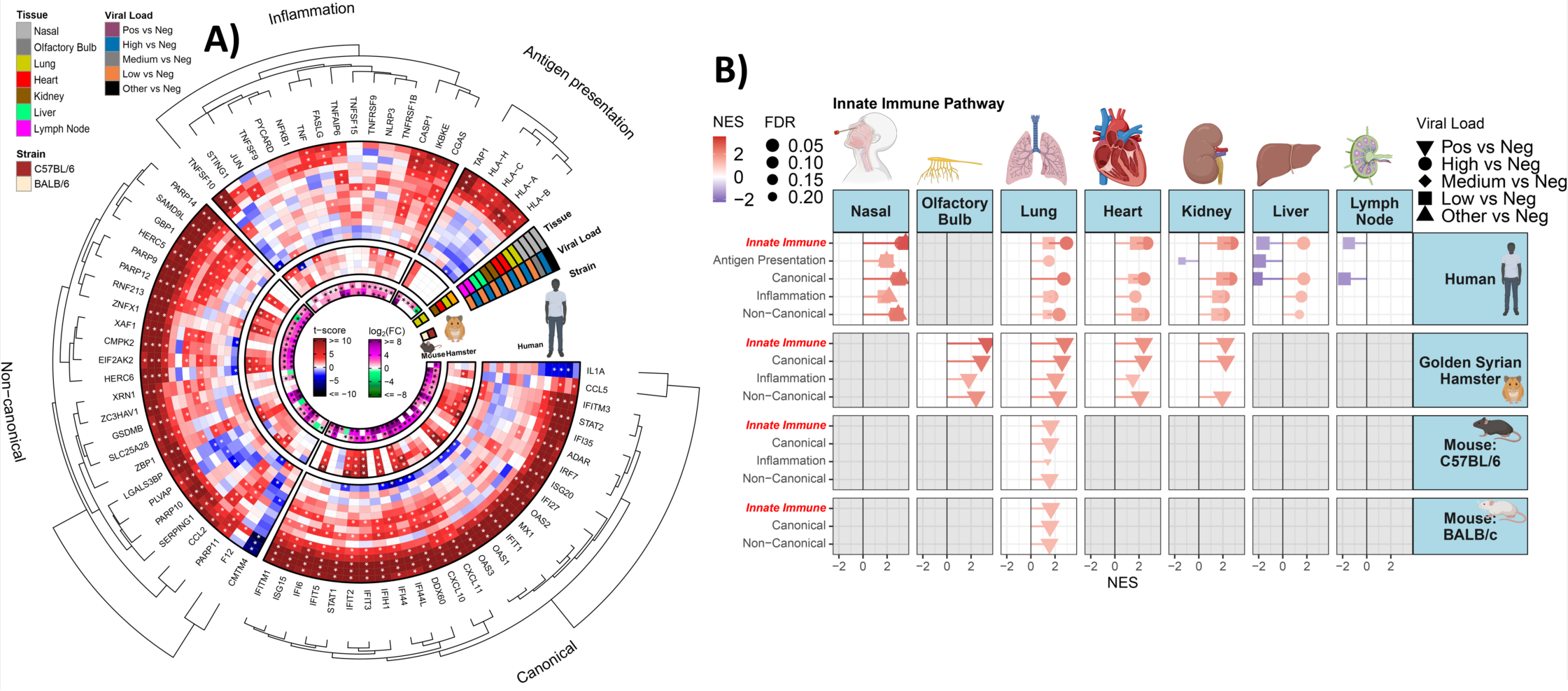
Mid- and late-stage and innate immune gene expression transcript changes in Infected Rodent Samples and human autopsy tissues from COVID-19 patients. **A)** Circular heatmap displaying the t-score statistics for specific innate immune genes, comparing SARS-CoV-2 infected hamster tissues, viral load versus negative patient nasopharyngeal and autopsy tissue samples, and log_2_-foldchange for SARS-CoV-2 infected C57BL/6 and BALB/c mouse lungs. **B)** Lollipop plots for custom gene sets determined by fGSEA (FDR <0.25), ranked by NES.

#### Non-canonical genes in autopsy tissues

These genes, introduced in the preceding section and all highly expressed in nasopharyngeal swab samples present a more heterogenous expression in autopsy tissue samples (**Fig. 2A**). These include: **1**) *GBP1*, an IFNγ, and cytokine signaling gene^43^, retaining high expression levels in the lung, heart, and kidney (**Fig. 2A**); **2**) Additionally, *HERC5* and *6* interact with canonical immune gene *ISG15* the former of which retains high expression in lung, heart, and kidney (**Fig. 2A**); **3**) The ISR initiating genes *EIF2AK2* and *MX1*^44^ are upregulated in the nasopharyngeal samples, but this augmentation is only retained in the kidney with significant down-regulation noted the in mediastinal lymph nodes. The upregulated *PARP9/12/14*, highly upregulated in the nasopharyngeal samples, remained upregulated in the autopsy lung, heart, and kidney (**Fig. 2A**). Additionally, *ZBP1*, *GSDMB*, *XAF1*, and *CMPK2* upregulated in the nasopharyngeal samples, retain their high expression in the lung, heart, and kidney and, to a lesser extent, in the liver.

#### Special implications of innate immune gene expression in mediastinal lymph nodes

Only 5 of 36 robustly upregulated innate immune genes in nasopharyngeal retain upregulation in mediastinal lymph nodes, while 15 are downregulated. Upregulated canonical IFN genes are *IFITM3, ISG15,* and *STAT2,* and non-canonical genes *RNF213*^45^ and DNA damage sensing genes, *cGAS* and *STING1.* The 15 reduced genes include 5 MHC1-related genes and also, *ZNFX1*, *SERPING1*, and *PVLAP* for which reduced expression can contribute to cell death/inflammation ^21,22,46^, and *HERC5*/*6* which upregulates *ISG15*^27^ (**Fig. 2A**). The critical implications of these expression patterns for compromised mediastinal lymph node immune function is addressed in more detail later.

#### Downregulated genes of interest in the autopsy tissues

Expanding our assessment of downregulated genes identified in the nasopharyngeal samples to human autopsy and animal model tissues revealed that these gene patterns were primarily relegated to nasopharyngeal samples (**Figs. S1A, S1B**) with several notable exceptions. Antimicrobial peptide/degranulation genes were downregulated in the lymph nodes and across both human and mouse lung samples and IL-1 family genes were downregulated in the hamster lung and heart samples (**Fig. S1B**. In total, our data showed that SARS-CoV-2 viral load has a negative association with genes in the broad classes of the IL-1 family, sulfonation/selenoamino acid metabolism, antimicrobial, and cornification/keratinization. The association of viral load and tissue site with IL-1 and degranulation highlights the importance of tissue-level specificity when assessing host responses to viral infection.

#### Gene expression in SARS-CoV-2 infected rodents

like our human tissues, lung, heart, and kidney of hamsters and lung of C57BL/6 and BALB/c mice show significant upregulation of most canonical and non-canonical innate immune genes. In the hamster brain, innate immune gene pathways were upregulated in the olfactory bulb and, to a lesser extent, in the cerebellum and striatum (**Figs. 2A, 2B, S2, S3**). Treatment of C57BL/6 and BALB/c mice with baricitinib or tofacitinib, which act as Janus kinase inhibitors, decreased activation of innate immune gene pathways (**Figs. S4, S5**). Together, these results demonstrate that innate immune genes are upregulated in the presence of replicating virus in early- and mid-infection in the human nasopharyngeal and rodent tissues and maintained despite the evidence of viral infection in other tissues.

**Fig 3.**
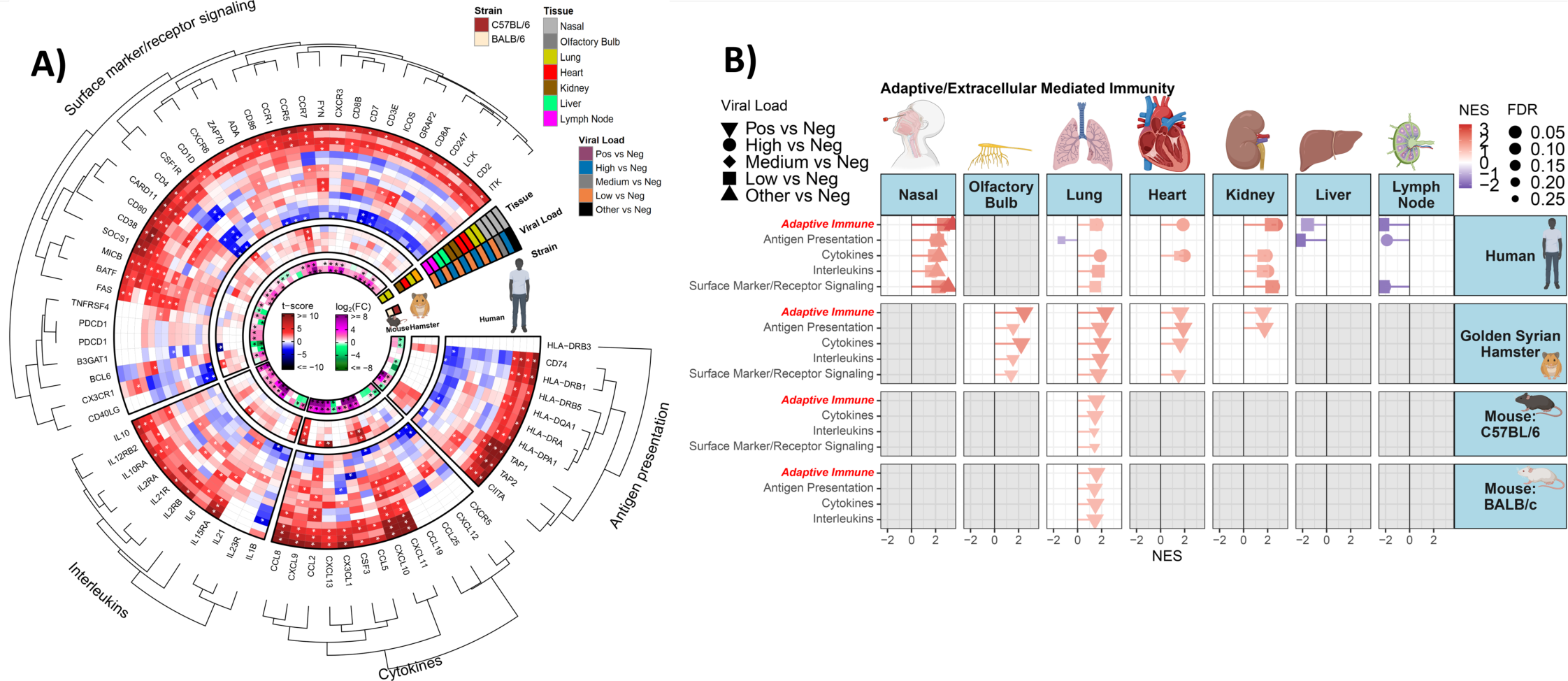
SARS-CoV-2 infection alters extracellular-mediated immunity-associated gene expression. **A)** Circular heatmap displaying the t-score statistics for extracellular-mediated immunity-associated genes comparing SARS-CoV-2 infected hamster tissues, viral load versus negative patient nasopharyngeal and autopsy tissue samples, and log2-foldchange for SARS-CoV-2 infected C57BL/6 and BALB/c mouse lungs. **B)** Lollipop plots for custom extracellular-mediated immunity-associated gene sets determined by fGSEA (FRD <0.25) ranked by NES.

#### Germline mutations in innate immune genes produce symptoms widely overlapping with those in severe COVID-19 patients

Considering syndromes induced by germline mutations in canonical and non-canonical genes (**Figs. S6, Table S3**) provides key clues to the importance of their dysregulation for lethal, inflammatory consequences in COVID-19. Such mutations of canonical immune genes produce Aicardi–Goutières syndromes (AGS) wherein overexpression of IFN-signaling genes results in constitutive cytokine-storm and often lethal outcomes in children^47^ (**Fig. S6, Table S3**). In rare families, loss-of-function germline mutations in *ZNFX1,* our top upregulated non-canonical gene in nasopharyngeal samples, produce often-lethal overaction of IFN-signaling only when affected individuals encounter pathogens^30^. *ZNFX1* is highly expressed in nasal swab samples but has little to no expression in all autopsy samples (**Figs. 2A, S6, Table S3**). As outlined in **Table S3**, germline mutations in other genes, including **1**) 14429G>A (p.Arg4810Lys) polymorphisms in the microvascular permeability regulator, *RNF213* lead to MOYA-MOYA disease generating intracranial artery atherosclerosis, systemic vascular diseases^45^. This gene, upregulated in nasopharyngeal samples trends downwards in the heart, kidney, and liver (**Fig. 2A**); **2**) germline loss-of-function mutations in the little-studied *PLVAP* gene, required for normal maintenance of microvascular permeability and gut barrier function, can cause death from severe diarrhea in infants^22^. *PLVAP,* while upregulated in nasopharyngeal samples is downregulated in heart, lung, liver, and kidney (**Fig. 2A**); **3**) germline loss and gain of mutations in, respectively two non-canonical innate immune genes, *SERPING1* and *F12*, which generate excess complement signaling, are leading causes of hereditary angioedema (HAE), a disorder associated with RAAS-overactivation^10^ Further importance of RAAS activation in our data is outlined later below. Patients with HAE have vascular and intestinal leakage and a propensity for thrombotic events^10,48^ as seen in lethal COVID-19^10,49^. Notably, *SERPING1,* while upregulated in nasopharyngeal samples, is downregulated in the lung, liver, and to a lesser extent kidney, while *F12* expression is increased in mediastinal lymph nodes (**Figs. 1, 2).**

### SARS-CoV-2 infection alters extracellular-mediated immunity-associated genes

In addition to the above transcriptional changes in innate and intracellular-mediated immunity genes, we also define the effects of SARS-CoV-2 infection on the dysregulation of extracellular-mediated immunity-associated genes. These queries include the categories of cell surface marker and surface receptor signaling, interleukins, cytokines, and class II antigen processing and presentation-related machinery (**Table S2**).

COVID-19 nasopharyngeal samples are highly enriched for upregulated innate (CSF1R, CCR1, CD80/86) and adaptive (CD3E/4/8/247) immune genes (**Figs. 3A, B**). In contrast, mediastinal lymph nodes, a central draining lymph node for the thoracic compartment, have marked downregulation of CD4- and CD8-associated genes inclusive of cell surface molecules, CD2/3E/4/8A/247 and T-cell receptor signaling (FYN, ITK, LCK). Interestingly, several innate and adaptive genes have differential expression across most tissue sites. MICB (NKG2D ligand), which functions as a stress-induced “kill-me” antigen, targeting expressing cells for clearance by natural killer cells^50^ is upregulated in most autopsy tissues. CSF1R and CD3E/4 have conserved downregulation across most tissue sites (**Fig. 3A**). CSF1R is the receptor for both CSF and IL-34 and is critical for directing differentiation and proliferation of mononuclear phagocytes, an essential class of highly professional antigen-presenting cells^51^ while CD3E/4, T lymphocyte helper T-cell genes play key roles in adaptive immune responses^52^. Importantly, expression of class II MHC antigen processing- and presentation-related genes is altered with their global upregulation in nasopharyngeal samples contrasting with downregulation in lymph nodes and a mixed expression pattern in other organ sites (**Fig. 3A**). Within these antigen-presentation-associated genes, CD74 warrants special mention for conserved downregulation across autopsy tissues (**Fig. 3A**) because of its functional importance for loading and chaperoning peptides for antigen presentation^53^.

In addition to the above-dysregulated tissue expression of immune cellular markers and antigen-presentation-associated genes (**Figs. 2A, 3A**), there are also altered expression patterns for interleukin and cytokine genes, thus suggesting the potential of differential lymphocytic and monocytic population recruitment ^54,55^. First, autopsy tissues have increased expression (**Fig. 3A, 3B**) of CXCL9/10/11, three major Th1-type chemokines which impart cellular chemotaxis through the CXCR3 receptor on both lymphocytic and monocytic populations and of CCL2/8, potent augmenters of monocytic cell type chemotaxis with the former having a role in severe SARS-CoV-2^56^. Also, there is a general increased expression of *IL15RA* across all tissue sites assayed, a potent lymphoid and myeloid cell activation inducer, which can facilitate an inflammatory milieu^57^.

In view of the tissue-specific expression of immune cellular markers and antigen-presentation-associated genes (**Figs. 2A, 3A**), we sought to define the expression profile of cytokine and interleukin genes, which are key regulators of immune cell localization and activation^42,56^. As anticipated, inflammatory interleukins and cytokine genes are upregulated in nasopharyngeal samples (**Fig. 3B**). The autopsy tissues also show a broad pattern of interleukin and cytokine augmentation, most manifest for CXCL9/10/11, CCL2/8, and IL15RA (**Fig. 3A**). CXCL9/10/11 are 3 major Th1-type chemokines, which impart cellular chemotaxis through the CXCR3 receptor on both lymphocytic and monocytic populations. CCL2/8 are potent augmenters of monocytic cell type chemotaxis, with CCL2 having a role in severe SARS-CoV-2^57^. In contrast to the cytokines, which are critical for cellular motility, IL15RA acts as a potent lymphoid and myeloid cell activation inducer, with the potential to facilitate an inflammatory milieu^58^. Together, these interleukin and cytokine patterns reveal a conserved potentiation of chemotactic gradients in human samples, which are known to attract both lymphocytic and monocytic populations in response to interferon and inflammatory stimuli.

Gene set enrichment analysis (GSEA) for the above-conserved patterns of gene perturbation, suggests significant directionality in human samples with a striking induction of extracellular mediated immunity gene sets for nasopharyngeal samples (**Fig. 3B**). Interestingly, in the kidney all gene sets except antigen processing and presentation are enriched, while lung and heart show augmentation of cytokine/interleukin or interleukin alone, respectively (**Fig. 3B**). Both liver and mediastinal lymph nodes show a conserved downregulation of extracellular mediated immunity, which is manifest for all gene sets in mediastinal lymph nodes, while liver has downregulated expression for all above gene sets, except cytokines (**Fig. 3B**).

Taken together, the above data reveal divergent transcriptional responses in lymphocyte-associated genes and class II antigen processing/presentation in SARS-CoV-2. Additionally, lymphoid organs from patients with severe SARS-CoV-2 present with transcriptional downregulation of adaptive immune-associated genes. These data are validated in our animal models of SARS-CoV-2 infection wherein extracellular mediated immunity genes are increased across all tissue sites in both Syrian hamster and mouse models of SARS-CoV-2 (**Fig. 3A, B**); further, the mouse model lung samples have a pattern more similar to human nasopharyngeal samples, while the Syrian hamster model demonstrates similarity to the human cytokine response (CCL2/8/10) across all assayed tissue sites.

### RAAS overactivation by upstream immune genes tracks with severe lethal COVID-19

RAAS signaling normally fluctuates to maintain blood pressure but overactivation induces severe inflammation as observed in HAE to trigger its overlapping features with lethal COVID-19^10^. As others have stressed, no single gene expression alteration causes pathogenic RAAS but rather the imbalanced patterns seen in our data below.

#### Imbalanced angiotensin axis

Chronic diminution of ACE2 and/or ACE levels and imbalanced angiotensin (AGT) regulatory axis induce overactivated RAAS and tissue-damaging events by altering AGT2 binding to AGTR1/2 receptors producing vasoconstriction, inflammation, fibrosis, tissue damage, edema, and activation of NADPH metabolism to increase NADPH to elevate intracellular ROS levels (**Figs. S6, Table S2)**. Mechanisms include activating renin (REN) renal secretion to generate AGT1 and cleavage of the latter by ACE1, or in the heart by CMA1 and to act through the MAS1 receptor to trigger the above inflammation consequences^10^ **Fig. 4A**, **S6**, **Table S2)**. In our autopsy samples, gene expression changes would favor *AGTR1* activation over *AGTR2/MAS1,* resulting in RAAS-activation as reflected by downregulation of *ACE2* and *AGTR2* in the lung; *CSTA* in the heart; and *MAS1* and, to a lesser extent, *AGTR2* in the lymph nodes (**Fig. 4B**).

**Fig 4.**
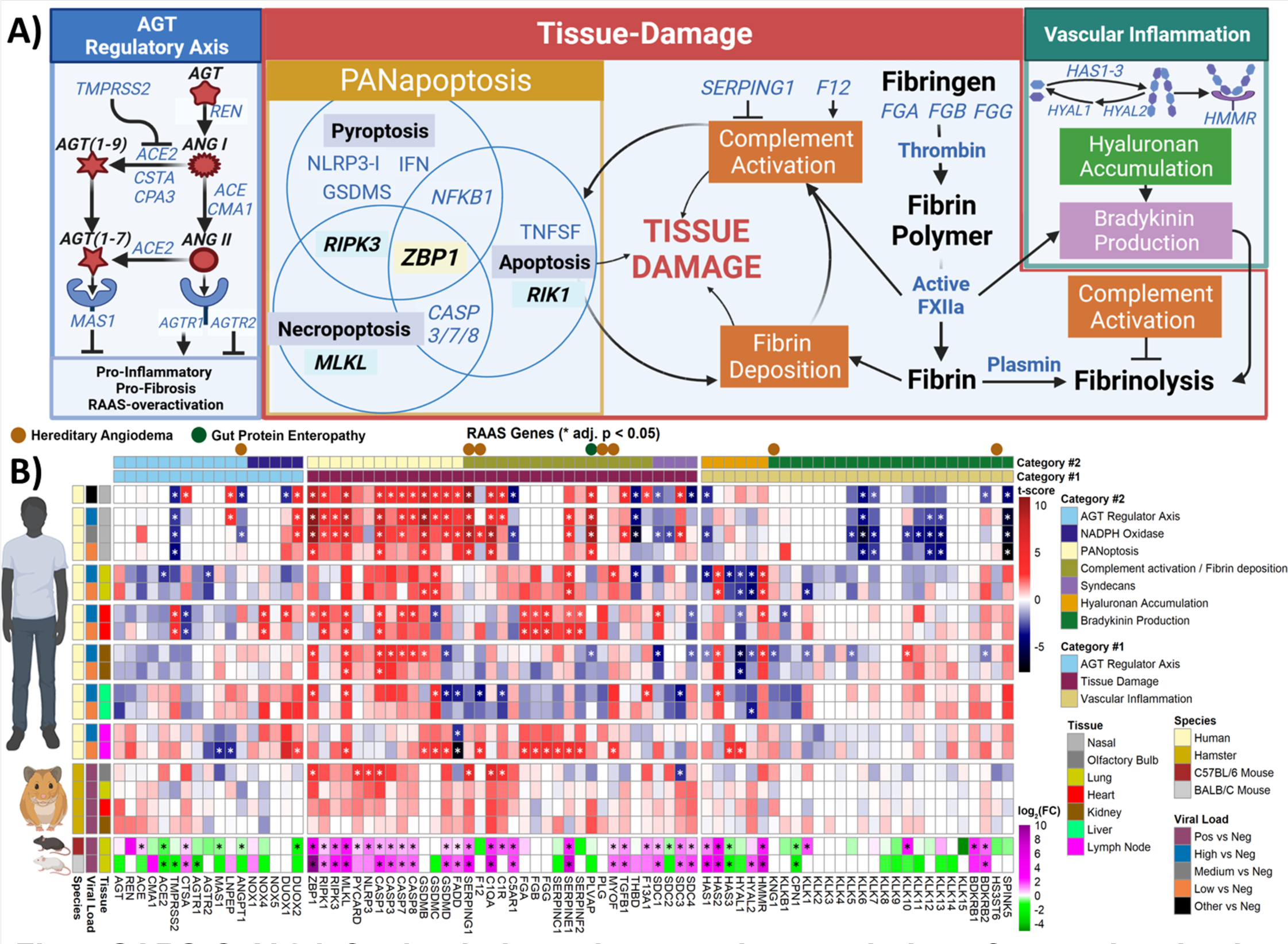
SARS-CoV-2-infection induces increased transcription of genes involved in RAAS-overactivation in the Human Lung, Heart, and Lymph. **A)** Schematic summary of RAAS pathway. **B)** Linear heatmap displaying the t-score statistics for RAAS target genes comparing SARS-CoV-2 infected hamster tissues, viral load versus negative patient nasopharyngeal and autopsy tissue samples, and log2-foldchange for SARS-CoV-2 infected C57BL/6 and BALB/c mouse lungs.

#### Overactivation of complement signaling

This key RAAS component, can be induced by the leading HAE germline loss and gain of function mutations respectively for two non-canonical innate immune genes, *SERPING1* and *F12*^10,21^. SERPING1 inhibits and F12 facilitates complement activation with accompanying altered downstream regulation of C1QA, C1R, and C1S genes^10,58,59^. Both *SERPING1* and *F12* are upregulated in nasopharyngeal samples but *SERPING1* is downregulated in the lung, liver, and to a lesser extent in the kidney, while *F12* is upregulated in the lymph nodes and somewhat in the heart (**Fig. 4B**). *C1QA* is upregulated in the nasopharyngeal, lung, heart, and lymph nodes, and *C1R* in the heart and lymph nodes while downregulated in the liver and *C5AR1*, is upregulated in the lung, heart, and lymph nodes (**Fig. 4B**). Additional germline mutations also induce HAE ^60^, including *KNG1*^61,62^*, PLG*^63,64^, *ANGPT1*^65^*, MYOF*^65^ (increased in lung and lymph nodes), and *HS3T6*^65^ (**Figs. 4B, S6, and Table S3**). All above gene changes can induce vascular disruption and collaborate with excess fibrosis damage as outlined further below.

**Fig 5.**
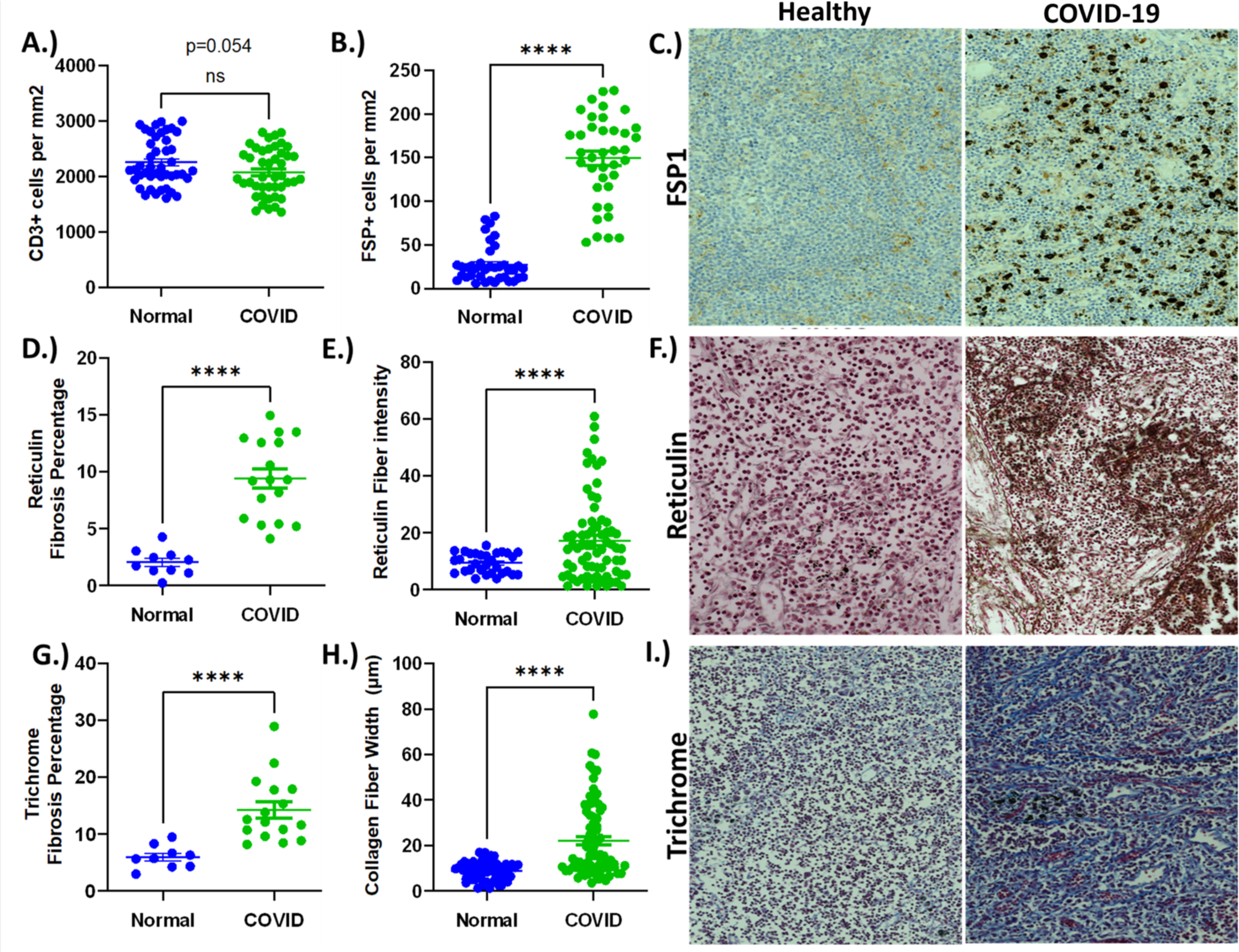
Severe COVID-19 disrupts organ architecture with an associated accumulation of pathogenic fibroblasts. Mediastinal lymph nodes were obtained from patients without lung disease or from patients who died from COVID-19. 5 regions (at 10X magnification were evaluated from each patient sample and evaluated for **A)** CD3 (cells/cm^2^) **B)** FSP (cells/cm^2^) **C)** Representative staining of FSP staining **D)** Reticulin staining and the percentage of area with reticulin staining **E)** Reticulin fiber intensity **F)** Representative staining of reticulin staining **G)** Percentage of fibrosis staining (based on Trichrome staining) **H)** Collagen Fiber Length **I)** Representative staining of Trichrome staining.

**Fig 6.**
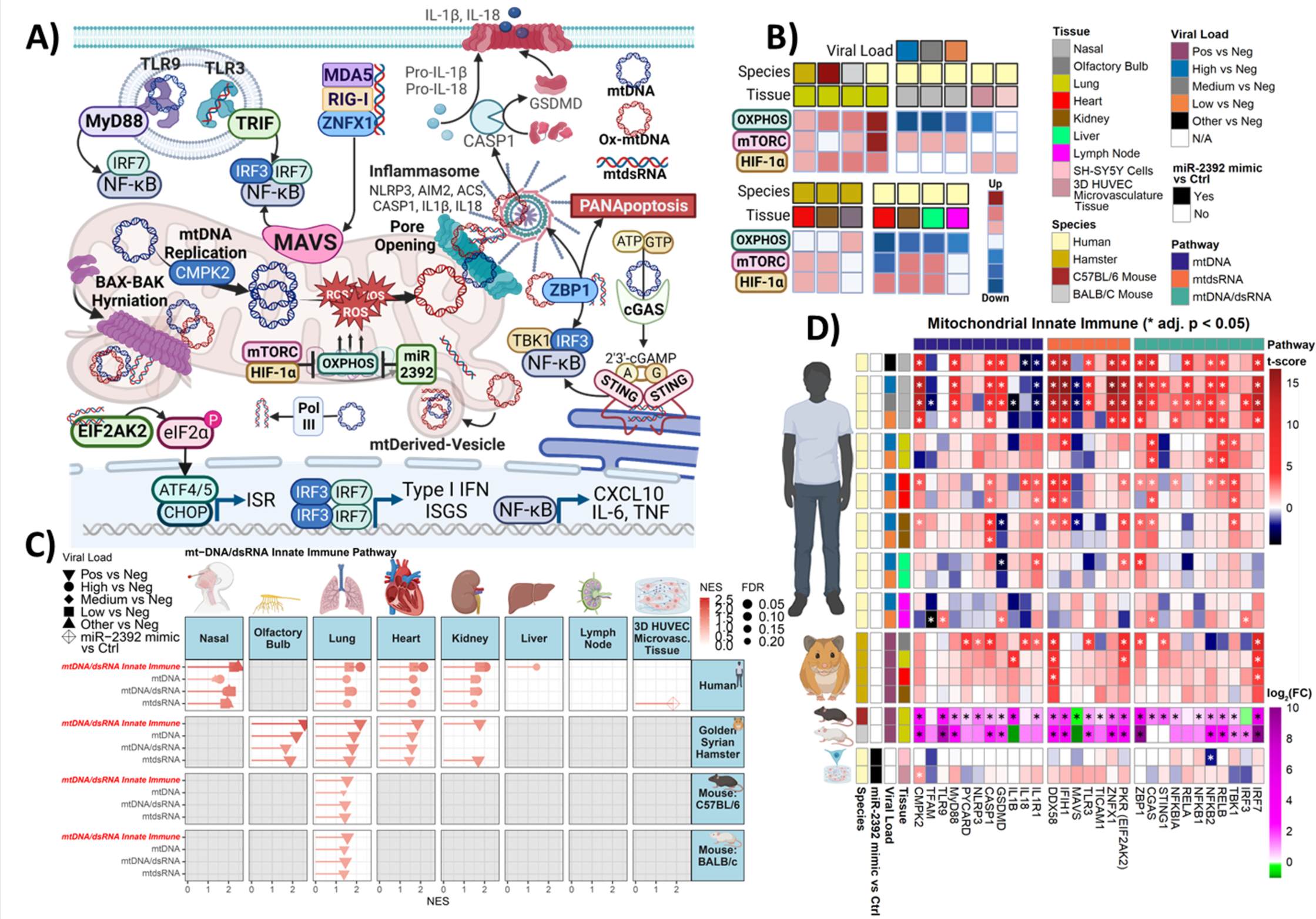
SARS-CoV-2-infection upregulates mtDNA/mtdsRNA-activated genes despite the absence of viral transcripts. **A)** Schematic summary of mtDNA/mtdsRNA-activated pathways. **B)** Graphical representation of findings from Guarnieri et al.^82^ **C)** Lollipop plots for statistically signifcant changes determined by fGSEA, ranked by NES, and **D)** Linear heatmaps displaying the t-score statistics for mtDNA/mtdsRNA-activated genes SARS-CoV-2 infected hamster tissues, viral load versus negative patient nasopharyngeal and autopsy tissue samples, miR-2392-expressing 3D-HUVEC-MT cells, and log_2_-foldchange (FC) for SARS-CoV-2 infected C57BL/6 and BALB/c mouse lungs.

#### PANoptosis

PANoptosis is driven by one of our, non-canonical immune genes, *ZBP1* to mediate multiple modes of cell death^32,55^. *ZBP1* is upregulated in nasopharyngeal samples (**Fig. 1 and 4B**) retains high expression in the lung, heart, and kidney and, to a lesser extent, in the liver (**Figs. 2A, 4B**). ZBP1 induces different PANoptosis death outcomes through differential binding to RIPK1, RIPK3, and MLKL^32,55^ as follows; binding to RIPK1 induces NF-κB-signaling and subsequent TNF and IL6 signaling^32,55^ and *TNF, ZBP1,* and *IL6* expression are coordinately increased in nasopharyngeal samples and to a lesser extent lung, heart, and kidney (**Fig. 4B**). Notably, IL-6, in conjunction with TGF, can also trigger the development of Th17 inflammatory cells. **2**) interaction with *RIPK3*, upregulated in heart and lymph nodes (**Figs. 4A, 4B**), can activate IFN and NF-κB, cGAS-STING, and mitochondrially-bound-NLRP3 inflammasome signaling, all driving DNA damage to induce dsRNA and/or dsDNA release into the cell cytosol which drives inflammasome signaling^32^. The RIPK3-ZBP1 interaction induces necroptosis through pyroptosis, a process mediated by gasdermin family proteins (GSDMs) which function with NLRP3, PYCARD, and caspases to induce swelling and rupture of cell and mitochondrial membranes^32,55^. *NLRP3* is highly expressed in the lung, kidney, and liver, *PYCARD* in the lung and kidney, and CASP1 in the nasopharyngeal, heart, kidney, and to a lesser extent in the lung and liver (***Figs. 4A, 4B***). *GSDM/B/C/D* are all significantly upregulated in nasopharyngeal samples and lymph nodes, *GSDMB/C* in the lung, kidney, and heart, and *GSDMD,* in the lung and heart (**Fig. 4B**). MLKL-ZBP3 interactions are also central to necroptosis ^32,55^ MLKL is significantly upregulated in nasopharyngeal, heart, kidney, liver, and lymph node samples (**Fig. 4B**). **5**) ZBP1-RIPK3 interaction with FAAD rather than MLKL switches cell death towards apoptosis associated with caspases 3/7/8^32,55^. *FAAD*, while highly expressed in nasopharyngeal samples, is significantly downregulated in the liver and lymph nodes, while *CASP3* is significantly upregulated in the kidney and highly expressed in all other sites except the lymph nodes*. CASP7/8* is significantly upregulated in the nasopharyngeal, heart, and kidney and to a lesser degree in the lung and liver samples (**Fig. 4B**) Thus, our overall data suggests ZBP1-RIPK3 interactions may drive necroptosis and some apoptosis-inducing PANoptosis in lethal COVID-19.

#### Fibrin Deposition

Excess fibrin deposition is a strong component of RAAS overactivation interacting with PANoptosis^10,58,59,66,67^. Despite no detectable expression in nasopharyngeal samples, three genes encoding fibrin synthesis enzymes, *FGA/B/G* are all significantly upregulated in heart and lymph nodes and, to a lesser extent, in lungs (**Fig. 4B**). Serpin protease family gene expression can initiate cycles of excess fibrin deposition turnover and another highly expressed, non-canonical immune gene in nasopharyngeal samples, *SERPINE1,* which encodes a tissue urokinase inhibits fibrinolysis^12,68–70^. This gene retains high expression in lung, heart, and lymph nodes, while *SERPINs C1* and *F2*, encoding plasma thrombin inhibitors^12^, while lacking expression in nasopharyngeal samples are significantly upregulated in lymph nodes and, to a lesser extent, lung and heart (**Fig. 4B**). Two other key RAAS over activation components, hyaluronic acid (HA) and bradykinin (BDK), interact with fibrin excess to cause vascular leakiness. HA binds to the HMMR receptor and increased HA in human plasma correlates with severe COVID-19, while blocking elevated HA in SARS-CoV-2 infected mice decreases mortality^12,71^. Three hyaluronan synthase isoenzyme genes, *HAS1/2/3*, *HAS2* are increased in the lung, liver, and kidney, and *HAS3* in lymph nodes. *HYAL2*, which degrades HA^72,73^, while having little expression in nasopharyngeal samples is significantly decreased in lung, kidney, and liver, while *HMMR* is significantly increased across lung, heart and liver samples (**Fig. 4B**). BDK production is facilitated by F12 to mediate conversion of pre-kallikreins KLK1-15 to kallikreins and their conversion to bradykinin by KNG1, KLKB1, and CPN1, which bind to BDKRB1/2^74^. SPINK5 is a serine protease that inhibits KLKS5/7/14. More nuanced changes in these genes occur in our data, but while having little or no expression in nasal swab samples, *KLK10* is significantly upregulated in the kidney, *KLK11* trends upwards in the heart, liver, and kidney, *KLK14* is upregulated in the kidney, liver, and lymph nodes and *SPINK5* is significantly downregulated in nasopharyngeal samples (**Fig. 4B**).

In conclusion, our data suggest that changes in the expression of upstream key immune genes track with being downstream drivers for RAAS overactivation contributing to PANoptosis, excess complement, fibrin, and HA deposition which induce vascular and other tissue damaging events in lethal COVID-19. Importantly, these changes suggest severe damage to mediastinal lymph nodes in lethal COVID-19 as explored more deeply just below.

### Severe SARS-CoV-2 disrupts lymphoid organ architecture with an associated accumulation of pathogenic fibroblasts

Laboratory changes, consisting of a battery of immunohistochemistry (IHC) studies validate the above transcriptomic changes further defining severe damage to mediastinal lymph nodes. First, as compared to autopsy controls there is a small reduction in CD3^+^ (**Fig. 5A**), which is consistent with the noted downregulation of immune cell markers (CD4, CD8A) (**Fig. 3A**). Second, our transcriptomic findings of increased expression of complement activation/fibrin deposition-related genes and association with excess fibrin deposition fit with the known recruitment of proliferative, collagen-producing fibroblasts to provide long-term remediation of tissue damage^75^. Indeed, there is a striking increase in the prevalence of a cellular population with IHC staining for fibroblast surface protein (FSP) (**Figs. 5B, C**). Third, in association with the above increase in fibroblasts, there is a marked increase in the lymph node samples for both fibrosis percentage (**Figs. 5D, F**) and fiber intensity (**Figs. 5E, F**) as assessed by reticulin staining. This histopathology stain assesses fiber deposition primarily composed of type III collagen and fibronectin and is consistent with fibrotic processes^76–78^. Importantly, reticulin fibrosis of lymph nodes has been previously reported to have an associated pathophysiology, in other disease settings, leading to impaired vaccination responses and cellular replenishment^79–81^. Fourth, as an extension to the above reticular fiber assessment, Mason’s trichrome staining for fibrosis reveals a pronounced increase relative to controls (**Figs. 5G, I**). Fifth, consistent with collagen being a major component of both reticulin and trichrome staining, there is an overall increase in collagen fiber width in the lymph node autopsy samples relative to controls (**Figs. 5H, I**). In summary, these IHC data reveal a reduction in CD3^+^ lymphocytes within the mediastinal lymph node co-occurring with large increases in fibrosis-associated markers (reticulin, trichrome, collagen), and recruitment of pathogenic fibroblasts.

### SARS-CoV-2 upregulates mtDNA/mtdsRNA-activated genes

Mitochondrial dysfunction activates the innate immune system via inhibition of OXPHOS and elevated mROS production, triggering the oxidation and release of mtDNA/mtdsRNA into the cytosol^13–17^. Like viral genomes, cytosolic mtDNA/mtdsRNA is detected by sensors of immunostimulatory RNA/DNA, leading to the induction of IFN, IRF, NF-κB, and their target genes, activating an innate immune response^6,13–17^(**Fig 6A**). In our last publication, analyzing the aforementioned rodent and human samples, we observed substantial alterations in mitochondrial transcripts and genes that regulate mitochondrial function, including HIF-1α, which upon activation, inhibits OXPHOS and mitochondrial biogenesis to favor glycolytic metabolism^82^ (**Figs. 6A, B**). Nasopharyngeal and autopsy heart, kidney, and liver samples displayed transcriptional inhibition of mitochondrial genes associated with OXPHOS and increased expression of HIF-1α-target genes (**Fig. 6B**). Lung autopsy samples and infected rodent tissues showed a general upregulation in mitochondrial gene expression and upregulation of HIF-1α-target genes. OXPHOS inhibition caused by decreased mitochondrial gene expression would increase ROS, increase HIF-1α activation, and trigger the release of mtDNA/mtdsRNA, activating an innate immune response.

#### Upregulated, mtDNA/mtdsRNA-activated genes in tissues with SARS-CoV-2 RNAs

In the human nasopharyngeal samples and rodent lungs, SARS-CoV-2 RNAs upregulated mtdsRNA-activated genes and genes involved in pathways activated by mtDNA including the cGAS-STING, NLRP3-I, and ZBP1 (**Fig. 6C**).

CMPK2 is the rate-limiting step for mtDNA replication, and the induction of mtDNA replication is central to the creation of oxidized mtDNA, which is released from the mitochondrion^13,38^. Cytosolic mtDNA binds/activates AIM2 and the NLRP3-I. Upon binding mtDNA, AIM2 and the NLRP3-I activate CASP1, triggering IL-1β/18 secretion and activating GSDMD, triggering pyroptosis^15,16^. Cytosolic mtDNA/mtdsRNA also binds/activates ZBP1, inducing PANoptosis and activating cGAS-STING^33–35^, and mtDNA binds TLR9, which with MyD88 activates IRF3 and NF-κB^17^. These aforementioned mtdsRNA-activated genes are all strongly upregulated in the nasopharyngeal samples, with the exception of *IL1β/18* (**Fig. 6D**). Additionally, in the rodent lungs, upregulated mtDNA-activated genes included *CMPK2*, *NLRP3, CASP1, PYCARD, ZBP1*, *TLR9*, and *AIM2* (**Fig. 6D**).

#### Upregulated, mtDNA/mtdsRNA-activated genes in the absence of viral transcripts

Despite significantly lower levels of SARS-CoV-2 transcripts (**Table S1)**, the human lung and human and hamster visceral tissues displayed significant upregulation of mtdsRNA-activated innate immune pathways After the nasal and mouse lung tissues, whose activation of mtDNA/mtdsRNA was aided by viral RNA, the human heart and lung displayed the next highest upregulation of these genes, followed next by human kidney and liver, and to a lesser extent, also hamster heart and kidney. Since there were no SARS-CoV-2 transcripts present in the COVID-19 autopsy samples tissue activation of these immunostimulatory sensors would have to come from the release of mtDNA/mtdsRNA.

The COVID-19 lung autopsy tissues exhibited a substantial upregulation of all the mtDNA/mtdsRNA-activated innate immune pathways (**Fig. 6C**). The human and hamster heart and kidney displayed strong upregulation of all mtDNA/mtdsRNA innate immune pathways, except for the unchanged NF-κB signaling in the human kidney (**Figs. 6C, D**). The inflammasome pathways were strongly upregulated in the liver, and consistent with the innate immune data, no pathways were significantly upregulated in the lymph nodes. For individual genes, the mtDNA cytosolic sensors *CMPK2, ZBP1, AIM2*, *CASP1*, *cGAS* and their downstream activators, as well as a majority of mtdsRNA sensor genes, were upregulated in the human heart, lung, kidney, and, to a lesser extent, in the liver (**Figs. 2B, 6D**). Hamster heart and kidney displayed an upregulation of *CMPK2*, *CASP1, GSDMD, STING, ZBP1*, and genes involved in NF-*κ*B-signaling (**Fig. 6D**).

#### *SARS-CoV-2* activates *the ISR* and unfolded protein response (*UPR*) *in several tissues*

UPR and ISR activation helps the cell adapt to environmental stressors and promotes cell survival; however, chronic signaling promotes inflammation and cell death (**Fig. S8**). Environmental stresses initiate the ISR through four specialized kinases that detect heme deprivation and OXPHOS inhibition via EFI2AK1/HRI, dsRNA via EFI2AK2/PKR, amino acid deprivation and oxidative stress via EFI2AK4/GCN2, and the accumulation of unfolded/misfolded proteins in the endoplasmic reticulum (UPR^ER^) via EFI2AK3/PERK^83,84^. EFI2AK1-4 activation triggers phosphorylation of the eIF2α that activates transcription factors ATF4/5 and their downstream target genes, including cytokines/death factors^84,85^ (**Fig. S8**). UPR^ER^ activation also upregulates genes involved in ER-homeostasis^85^, and the mitochondria UPR (UPR^MT^) upregulates antioxidant and mitochondrial maintenance genes^86,87^.

The ISR was significantly activated in all the human autopsy tissues except the lymph nodes (**Fig. S9A, B)**. Activation was highest in the lung and nasal tissues, followed by the heart, kidney, and to a lesser extent, the liver. The UPR^MT/ER^ was upregulated in the human lung, and the UPR^MT^ was downregulated in the heart and upregulated in the lymph nodes (**Fig. S10A, B)**. In rodents, the ISR and UPR^ER^ were upregulated in the mouse lungs, and the ISR, to a lesser extent, in the hamster lung, kidney, and heart (**Figs. S10, 11)**. Consistent with chronic stress signaling, the ISR and UPR were most upregulated in the autopsy lung samples, followed by the mouse and hamster lung samples. For the visceral tissues, the human heart and kidney displayed increased transcription of ISR-target genes compared to their hamster counterparts, suggestive of increased ISR activation with COVID-19 progression in these tissues.

For individual genes, the dsRNA sensing *EFI2AK2/PKR* was upregulated in all the tissues except for the lymph nodes, for which it was downregulated (**Fig. 2B, S9B)**. *EFI2AK4/GCN2* or *EFI2AK1/HRI*, which sense mitochondrial dysfunction, were upregulated in the human tissues (**Fig S9B)**. *PGC-1α*, which regulates mitochondrial biogenesis and quality control^88^, and *GPX4*, *AIFM2*, *PARL*, and *STARD7*, which detoxify lipid-peroxidation and inhibit ferroptosis^89,90^, were strongly downregulated in the human heart and kidney. Their downregulation would elevate mROS and the innate immune response in these tissues (**Fig. S10A)**.

#### miR-2392 inhibits OXPHOS and upregulates ISR and mtDNA/mtdsRNA-activated genes in the absence of viral transcripts

SARS-CoV-2 infection results in the activation of micro-RNA 2392 (miR-2392), and treatment with anti-miR-2392 therapeutics antagonizes SARS-CoV-2 infection in cells and hamsters^91^. MiR-2392 contains seed sites for 362 nDNA OXPHOS genes, and expression of a miR-2392 mimic in 3D-HUVEC-MT cells resulted in a robust downregulation of nDNA OXPHOS genes^82^ (**Fig. 6A, B**). Expression of miR-2392 significantly upregulated all the innate immune pathways (**Fig. S11A, S11B)**, including the ISR and mtDNA/mtdsRNA-activated pathways and *CMPK2 (***Fig. 6C**), indicating that miR-2392 can contribute to the overactivation of the innate immune response through OXPHOS inhibition. miR-2392-expressing 3D-HUVEC-MT cells displayed increased expression of cytokine/chemokine ISR-target genes and significant downregulation of antioxidant and autophagic ISR-target genes and genes of the UPR^ER^. This could contribute to miR-2392-induced mitochondrial dysfunction, augmenting viral replication and the innate immune response (**Figs. S10**, **S11)**.

Despite the absence of viral transcripts, we observed significant upregulation of mtDNA/mtdsRNA-activated genes that correlated with the inhibited transcription of OXHPOS genes, which was most pronounced in heart and kidney autopsy samples (**Fig. 6**). Furthermore, we discerned a robust activation of HIF-1α, which was most evident in heart and lung autopsy samples (**Fig. 6**). These data and associated observations are consistent with the hypothesis that SARS-CoV-2-induced mitochondrial dysfunction can contribute to activation of the innate immune response through eliciting miR-2392 and the release of mtDNA/mtdsRNA into the cytosol. In all tissues, we also observed increased upregulation of ISR genes, which, when chronically activated, are a major factor in promoting inflammation and cell death (**Fig. S9)**.

### PBMCs and whole blood samples from early and later variants of SARS-CoV-2 display comparable immune responses

Peripheral blood mononuclear cells (PBMCs) from patients infected with Alpha and Omicron strains, as well as, whole blood from patients infected with early and later SARS-CoV-2 strains were analyzed (see Tables S6 and S7 for patient genotype and demographic summaries for whole blood samples). These peripheral compartment-derived samples display robust upregulation of the canonical innate immune, RAAS, and mtDNA/mtdsRNA-activated innate immune genes, comparable to the human nasopharyngeal samples (**Figs. 7A, S12**). Additionally, 18 of the 21 non-canonical immune genes upregulated in the SARS-CoV-2 positive nasopharyngeal samples, as well as genes involved in the inflammasome and NF-κB, are also significantly upregulated in the PBMCs and blood samples collected from SAR-CoV-2 infected individuals **(Fig. S12)**. Eight genes show consistent upregulation in our SARS-CoV-2 positive tissues and PBMCs and blood samples collected from Alpha- and Omicron-infected individuals (**Fig. 7B**). These genes are five mitochondrial-associated innate immune genes, *CMPK2, CGAS, ZBP1, EIF2AK2,* and *GDF15.* Also elevated are *MICB*, which encodes an antigen for clearance by natural killer cells^50^; and cytokines *CXCL10* and *CXCL11*, associated with COVID-19 severity^41^ (**Fig. 7B**).

**Fig 7.**
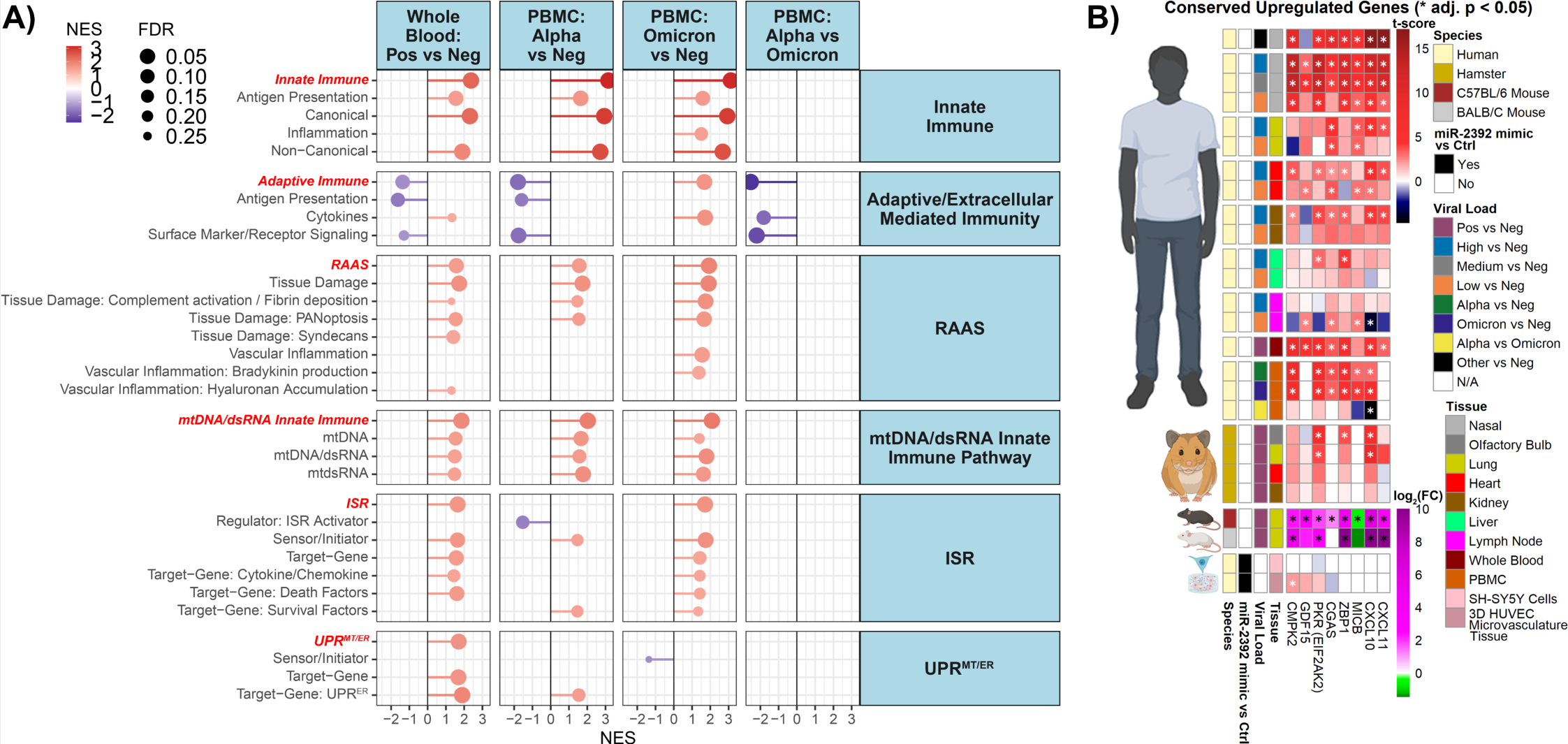
Transcript changes in PBMCs collected from SARS-CoV-2-Alpha ans SARS-CoV-2-Omicron-infected patients display comparable immune responses. Lollipop plots for statistically significant changes in custom gene sets determined by fGSEA for Whole blood and PBMCs ranked by NES. **B)** Linear heatmap displaying the t-score statistics for conserved upregulated genes comparing PBMCs collected from SARS-CoV-2-Alpha and SARS-CoV-2-Omicron-infected patients versus non-infected patients.

PBMC and blood from patients infected with the Alpha variant display a robust downregulation of genes involved in the adaptive and extracellular mediated immunity (**Fig. 7**), which generally recapitulates the response of the autopsy lymph nodes (**Figs. 3A, SF12**). In PBMCs, genes involved in class II antigen processing and presentation (*HLA-DRA, HLA-DRB1, HLA-DQA1*), TCR signaling (*LCK, FYN*), and lymphocyte state (*CD8A, CD8B, CD4, CD3E,* and *CCR7*) are significantly downregulated in Alpha-infection relative to healthy controls (**Fig. S12).** In contrast, the Omicron infection is associated with an overall muted transcriptional response, with the notable exception of monocytic chemotaxis (*CCL2, CCL8*) augmentation (**Fig. S12)**. These data reveal strain-specific behavior in the peripheral compartment, with the caveat that we cannot define whether these are based on the virus alone or a host viral interaction mediated by prior vaccination or infection. In aggregate, genes of innate immune, RAAS, and mtDNA/mtdsRNA in PBMCs are similarly upregulated in Alpha and Omicron variant infections; however, extracellular mediated immunity-related genes show divergent expression patterns wherein Alpha downregulates while Omicron upregulates these genes.

## DISCUSSION

Our present study offers novel insights that increase understanding of the inflammation dynamics that drive lethal outcomes for patients infected by SARS-CoV-2. *First*, we expand the dimensions of cytokine storm by linking alterations in canonical with an upstream role for “non-canonical” IFN-signaling genes to induce signaling overactivity of the RAAS pathway. Our study is informed by evaluating the effects of several genes for which germline mutations cause HAE, an inflammatory syndrome with tissue damage widely overlapping that seen in severe COVID-19^10,48^. Especially revealing are dynamics for germline loss and gain of-function mutations respectively for *SERPING1* and *F12*, each resulting in excess complement activity to induce RAAS overactivation components increasing fibrin deposition. These findings point to the mechanism by which COVID-19 induces systemic vascular leakage, loss of gut wall integrity, and thrombosis. Findings for rare germline mutations in genes such as *PVLAP*, which produces a similar loss of vascular and gut integrity, cause a similar transcriptional profile as COVID-19. Also, we reveal a role in in COVID-19 infection for *ZNFX1*, a little-studied gene for which rare germline mutations render patients incapable of mounting a response to pathogens. For RAAS overactivation, we emphasize, as have others^48^, that changes for expression of any one gene alone cannot be relied upon to judge overall or even a component of RAAS overaction. Rather, an integration of many genes and their ramifications for pathway signaling must be utilized to demonstrate an imbalance in gene expression driving abnormal RAAS activation.

*Second*, our study brings into focus important roles for multiple proteins encoded that shuttle from the nucleus to mitochondria during viral infection to foster activation of inflammasome signaling^6,13–17^. Importantly, this signaling occurs when the mitochondrion is damaged by mROS accumulation. Key proteins include *ZNFX1, CMPK2,* and our non-canonical innate immune genes functioning as RAAS components, including *GSDMB, XAF1,* and *ZBP1*. The roles of these latter three genes in inducing multiple modes of cell death likely play an integral part in lethal COVID-19. These data link response dynamics of the host to RAAS-overactivation and the mitochondrial ISR, to inflammation and cell death. Changes in ISR gene expression are linked to mitochondrial dysfunction through specialized kinases including EFI2AK1/HRI which detects the UPR^MT^ via OMA1 activation and DELE1 cleavage, EFI2AK2/PKR which detects dsRNA, and EFI2AK4/GCN2 which senses amino acid deprivation and oxidative stress^83,84^. OXPHOS inhibition activates HIF-1α via elevated mROS production, triggering the oxidation and release of mtDNA/mtdsRNA into the cytosol, and activating immunostimulatory dsRNA/dsDNA sensors.

*Third*, the clinical significance of our transcriptomic findings in mediastinal lymph nodes is confirmed by demonstrating excess fibrosis and collagen deposition accompanying a fundamental shift in their architecture and lymphocyte compartmentalization (**Fig 5**). General lymphoid organ dysregulation in severe COVID-19 has been described previously in reports that define lymphocyte depletion broadly^92–94^, including T-follicular cell helper deficiencies^95,96^, fibrin deposition^97^, and germinal center disruption^97–100^ further elucidating the basis of COVID-19 lymphopenia (**Fig. 3**). Activation of the acute fibrin response associated with fibrosis in the lymphoid organs may be analogous to the long-term dynamics in wound healing^78^, which we correlate with overactivated RAAS signaling. Hence, damage to the mediastinal lymph nodes could be critical for the defective adaptive immune response that severely comprises immune function leading to the lethal outcomes in COVID-19. *Finally*, our data may shed light on the persistent, unresolved lymphoid organ dysfunction observed in PASC or “long COVID”. For example, a recent study that tracked patients with PASC found dysfunctional B- and T-cell responses with elevated Type I IFN responses still present at 8 months post-acute disease resolution^101^.

While the innate and adaptive immune responses to SARS-CoV-2 are diverse and complex, there is increasing evidence of the integration between various cellular compartments in mounting a unified defense against SARS-CoV-2 infection. We have found that SARS-CoV-2 infection acts to inhibit mitochondrial proteins that block transcription of mitochondrial genes encoded in both the nuclear and mitochondrial DNAs^82^. This mitochondrial inhibition stimulates the induction of the non-canonical innate immune, *CMPK2*, which regulates the rate-limiting step of mtDNA replication, which in conjunction with mROS, results in the release of mtDNA through the mitochondrial permeability pore to activate the inflammasome and cGAS-STING pathways^16^. This linkage between mitochondrial dysfunction and the innate immune system links to these dynamics and SARS-CoV-2 induction of ZNFX1 and ZBP1. The SARS-CoV-2 hyper-induction of the ZNFX1 protein activates the interferon response by interacting with MAVS^31^ and the ZBP1 protein binds the telomeric-repeat-containing RNAs (TERRA) and the TERRA-bound ZBP1 oligomerizes into filaments on the outer mitochondrial membrane to activate MAVS^35^. Thus, the innate immune system links nuclear telomeres with cytosolic nucleic acid receptors and mitochondrial nucleic acid signaling.

In summary, we have defined unifying gene expression dynamics of innate and adaptive immunity that results from SARS-CoV-2 infection. These transcriptional changes have revealed that overactivation of RAAS signaling and mitochondrially mediated inflammation activation holistically comprise the cytokine storm that causes lethal COVID-19.

### Limitations of the Study

This study has several limitations. First, some of the clinical components of the study were designed retrospectively. Consequently, COVID-19 and control groups are not matched for demographic variables, comorbidities, or in-hospital treatments. Second, the transcriptomic results, alone must rely on correlational interpretation without demonstrating causation. However, orthogonal data derived from rodent SARS-CoV-2 infection models provide a measure of validation for key findings uncovered by these correlational studies. Furthermore, quantitative immunohistochemistry analyses confirm cellularity and architecture changes, which are directly correlated with our transcriptomic data. Certainly, the key conclusions we draw both require and invite validation in follow-up studies in larger cohorts.

## MATERIALS AND METHODS

**Study Design** To determine the effects of SARS-CoV-2 infection on mitochondrial function in patients, we calculated the relative expression levels of host genes in RNA-seq data from 735 nasopharyngeal samples and 40 autopsy cases from SARS-CoV-2 positive and negative individuals. We then compared the gene expression levels of infected versus control subjects using our curated set of immune gene lists (**Table S2).** Hamster^4^, mouse^102^ and PBMCs from COVID-19 patients infected with the Alpha^103^ or Omicron^104^ strains studies extended the conclusions. Whole blood RNAseq data was also derived from PAXgene specimens collected from 221 Military Health System beneficiaries with SARS-CoV-2 infection who enrolled in the Epidemiology, Immunology, and Clinical Characteristics of Emerging Infectious Diseases with Pandemic Potential (EPICC) study^105^. All animal experiments were conducted using approved standard operating procedures and safety conditions for SARS-CoV-2 in BSL3 facilities designed following the safety requirements outlined in the Biosafety in Microbiological and Biomedical Laboratories, 6th edition with protocols approved by the Institutional Animal Care and Use Committee (IACUC) at Icahn School of Medicine, Mount Sinai, NY^106^ and The University of North Carolina (UNC), Chapel Hill^107^. For all experiments, the preprocessing was blinded until the higher-order analysis which is described below involved determining fold-change values, t-scores, pathway analysis, and statistics. No outliers or data was excluded from any of the analysis.

### Human autopsy and nasopharyngeal samples collection and RNA Sequencing and Analysis

Nasopharyngeal specimens were collected from 216 SARS-CoV-2 -positive patients and 519 SARS-CoV-2 -negative patients, and autopsy heart, kidney, liver, lymph node, and lung samples were collected from 35-36 patients and 5-8 controls. Tissue samples were provided by the Weill Cornell Medicine Department of Pathology. The Tissue Procurement Facility operates under Institutional Review Board (IRB) approved protocol and follows guidelines set by the Health Insurance Portability and Accountability Act. Experiments using samples from human subjects were conducted in accordance with local regulations and with the approval of the IRB at the Weill Cornell Medicine. The autopsy samples were collected under protocol 20–04021814. Autopsy consent was obtained from the families of the patients. The subdivision of autopsy subjects into “high viral load” and “low viral load” on “admission” was determined from the nasopharyngeal swab samples from COVID-19 patients at the time of infection and hospital visit. The viral loads were assessed with a qRT-PCR cycle threshold (Ct) value of SARS-CoV-2 primers, with Ct of less than or equal to 18 being assigned to “high-viral load” label and Ct between 18 and 40 being assigned to “low viral load” classes. Ct values above 40 were classified as negative. The generation and processing of these tissues and RNA sequencing have been previously reported in Guarnieri et al,^82^. RNAseq data was processed as described by Butler et al.^7^ and Park et al.^108^. DESeq2^109^ was utilized to generate the differential gene expression data.

### Collection and RNA sequencing of whole blood samples from the EPICC Cohort

Whole blood specimens (PAXgene tube) were collected from the Epidemiology, Immunology, and Clinical Characteristics of Emerging Infectious Diseases with Pandemic Potential (EPICC) study. The full detail of this study is described in^105^. Briefly, the EPICC cohort study enrolls U.S. Military Health System (MHS) beneficiaries; eligibility criteria include a history of SARS-CoV-2 infection. Study procedures include collection of demographic and comorbidity data, COVID-19 vaccination data, acute COVID-19 illness clinical data, blood specimens, and other biospecimens. The EPICC study was approved by the Uniformed Services University Institutional Review Board (IDCRP-085) and all study participants provided consent when enrolled in the study. This study was conducted following good clinical practice and according to the Declaration of Helsinki guidelines.

Whole blood used for these RNAseq analyses was collected from 221 participants enrolled during 2020 and 2021 at eight Military Treatment Facilities (MTFs) in the United States: Alexander T. Augusta Military Medical Center, Brooke Army Medical Center, Madigan Army Medical Center, Naval Medical Center Portsmouth, Naval Medical Center San Diego, Tripler Army Medical Center, Walter Reed National Military Medical Center, and the William Beaumont Army Medical Center^105^.

Whole blood transcriptomic data was derived from collected whole blood PAXgene specimens as follows: Whole blood-derived total RNA was used as input at 200-600 ng for library preparation using the TruSeq Stranded Total RNA with Ribo-Zero Globin and IDT for Illumina - TruSeq RNA UD Indexes (96 indexes, 96 samples). Sequencing libraries were evaluated for size distribution using an Agilent Fragment Analyzer and for yield by qPCR using the Roche Light Cycler II 480 and Kapa Library Quantification Kit for Illumina Platforms. Sequencing libraries were pooled and used as input for sequencing on the NovaSeq 6000 Platform on a NovaSeq 6000 S4 Reagent Kit v1.5 (200 cycles). Raw sequencing data was demultiplexed to FASTQ and processed for mapping and read counts by STAR within the seqEngine workflow. Differential Gene Expression (DGE) was determined using DESeq2 with a filter for genes with at least 10 or higher counts per million (CPM) in at least 10 samples. The fold-change, t-score, and p-adjusted values were utilized from the DESeq2 output for all downstream analyses.

### SARS-CoV-2 model and RNA-sequencing of hamster samples

The hamsters were infected by intranasal inoculation with SARS-CoV-2WA, and lung, kidney, heart, and brain (olfactory bulb, striatum, cerebellum) samples were collected at the maximum lung viral titer 3 DPI. To approximate likely human nasopharyngeal infection progression, we analyzed gene expression profiles in the hamster tissues at 3DPI. The generation and processing of hamster tissue and RNA samples were performed as described in Frere et al,^4^.

### SARS-CoV-2 model, and RNA-sequencing of C57BL/6 and BALB/c mice treated with or without baricitinib or tofacitinib

Mice were housed in the UNC ABSL3 facility on a 12:12 light cycle using autoclaved cages (Tecniplast, EM500), irradiated Bed-o-Cob (ScottPharma, Bed-o-Cob 4RB), ad libitum irradiated chow (LabDiet, PicoLab Select Rodent 50 IF/6F 5V5R), and autoclaved water bottles. Animals used in this study included female 16-week-old C57BL/6J (B6) (The Jackson Laboratory stock 000664) or 10-12-week-old BALB/cAnNHsd (BALB/c) (Envigo order code 047) mice, purchased directly from vendors.

Mice were infected following light sedation, using 50 mg/kg ketamine and 15 mg/kg xylazine, by intranasal inoculation with 10^4^ pfu SARS-CoV-2 MA10 diluted in 50 μL PBS or PBS alone (mock infection). Blinded treatment groups for mice were used throughout the study to limit investigator subjectivity. 24 hours post infection mice were dosed with baricitinib 10 mg/kg or tofacitinib 50 mg/kg. 7 days post-infection mice were euthanized by an overdose of isoflurane anesthetic and lung tissues were collected for subsequent processing.

BALB/c or C57BL/6 mice were infected by intranasal inoculation with 10^4^ pfu SARS-CoV-2-MA10, and lung samples were collected at 4 DPI, after viral loads had peaked and declined. The generation and processing of the BALB/c or C57BL/6 lung tissue samples was performed as described in Guarnieri et al,^82^. BALB/c or C57BL/6 mice were infected by intranasal inoculation with 10^4^ pfu SARS-CoV-2-MA10, and lung samples were collected at 4 DPI, after viral loads had peaked and declined. To approximate midway between the human nasopharyngeal and autopsy viral titers, we analyzed gene expression profiles in hamster tissues at 4 DPI.

RNA from inferior mouse lung lobes was extracted using TRIzol Reagent (ThermoFisher Scientific), followed by overnight precipitation at −20°C, and quantified using a NanoDrop spectrophotometer (ThermoFisher Scientific). Ribosomal RNA from 1000 ng total extracted RNA was depleted using a NEBNext rRNA Depletion Kit (Human/Mouse/Rat; New England Biolabs Inc.). The remaining RNA was used to produce the sequencing libraries using the NEBNext Ultra II Directional RNA Library Prep Kit for Illumina (New England Biolabs Inc.) with AMPure XP (Beckman Coulter Life Sciences) for all bead cleanup steps. The libraries were sequenced on a NovaSeq 6000 System, using a NovaSeq 6000 SP Reagent Kit v1.5 (Illumina).

### Transfection of 3D HUVEC tissue and SH-SY5Y with miR-2392 mimics

Mature human 3D microvascular tissue models were grown by seeding human umbilical vein endothelial cells (HUVECs) (Lonza Cat# CC-2519) into a collagen/matrigel mixture as described^82,91^. Briefly, HUVECs were first grown in EGM Endothelial Cell Growth Medium (Cat# CC-3124), then cells (1 x 106/ml) were embedded into a collagen/matrigel mixture (Cat# 354236 and Cat# 356230). Microvessels were grown in EGM-2 Endothelial Cell Growth Medium (Cat# CC-3162) plus 50 nM Phorbol-12-myristate-13-acetate: (PMA) (Cat# 524400) for 7 days to form tubular microvessels before irradiation.

These tissue models were used to study vascular changes associated with miR-2392 mimics. The 3D HUVEC tissues were incubated according to the manufacturer’s instructions with the miR-2392 mimics or control lentivirus particles (MOI 1) for 48 hours. Specifically, we used the shMIMIC Lentiviral microRNA hsa-miR-2392 hCMV-TurboGFP UHT kit with the SMARTvector Non-targeting hCMV-TurboGFP Control Particles for the control vehicle transfection (Horizon Discovery Biosciences Limited, Cat #: VSH7357 and VSC10236). The tissue constructs used for miR-2392 mimics were dissolved in TRIzol (ThermoFisher Cat# 15596026) 48 hours post-transfection, without any fixation or homogenization.

Cell culture, RNA analysis, and data processing of miR-2392 mimic experiments in SH-SY5Y cells and RNA-seq were conducted as described previously^91^.

### RNA-seq of 3D HUVEC and SH-SY5Y tissue transfected with miR-2392 mimics

Details for RNA-seq and pre-processing were previously described^82,91^. All other analysis is similar to what is described for the other RNA-seq data present in this manuscript.

### Analysis combining sample RNA-seq data

To combine the results from the autopsy, nasopharyngeal swab, hamster, mice, PBMCS, miR-2392 mimics, and whole blood RNA-seq data, we utilized the t-score values from the DESeq2 analysis. Heatmaps were displayed using pheatmap (version 1.0.12). Circular heatmaps were produced in R (version 4.1.0) using the Complex Heatmap (version 2.9.4) and circlize (version 0.4.12) packages.

### Gene Set Enrichment Analysis (GSEA)

For pathway analysis, we utilized fast Gene Set Enrichment Analysis (fGSEA)^110^. Pathway analysis was done utilizing custom-made Gene Set files (available in **Table S2**). Using fGSEA, all samples were compared to controls, and the ranked list of genes was defined by the t-score statistics. The statistical significance was determined by 1000 permutations of the gene sets and a cutoff of FDR < 0.25 was used throughout the paper as recommended by GSEA instructions^111^. Lollipop plots were made using R (version 4.2.1) using ggplot2 package (ver 3.4.2).

### Mediastinal Lymph Node Histopathology

Lymph nodes were fixed via 10% neutral buffered formalin inflation, sectioned, and fixed for 24 hours before processing and embedding into paraffin blocks. Freshly cut 5-μm sections were mounted onto charged slides. Slides were stained using FSP1 antibody, hematoxylin, and eosin, reticulin, and Masson’s Trichrome according to standard protocol. Four 20× regions were randomly selected from each slide the Color deconvolution2 algorithm for ImageJ and cellular trichrome-rich areas were manually selected and measured for pixel area (cellular areas on red deconvolution, trichome on blue deconvolution). The total image pixel area was used to determine the percent fibroblast-rich trichrome-positive zones.

### Data Analysis Methods for Figures 1A and 1B

All analyses were performed in R v4.3.1.

#### 1. Regression Analysis on Viral Load

**Sample Filtering**: Samples were excluded from analysis if their total human sequence read count was less than 10e06. Number of subjects passing requirements: 486. **Viral gene count background estimation**: Viral sequence counts were found to have a direct relationship with PCR levels. From that analysis, we determined that the background (false positive rate) of SARS-CoV-2 virus sequences in non-infected individuals was approximately 100 reads per sample. We replaced all ‘0’ reads in our viral sample counts from uninfected individuals with a random integer between 1-100. Viral count data was log-transformed. **DE analysis**: We model human gene expression counts vs. log-transformed SARS-CoV-2 gene expression counts as described above. We first filtered for all human genes with at least 100 reads total across the 486 samples. Analyses were performed in R v4.3.1. Using Bioconductor packages DESeq2 v1.36.0^109^, sva v3.44.0 (https://doi.org/doi:10.18129/B9.bioc.sva) and apeglm v 1.18.0^112^, we fit a model of viral sequence counts vs. human gene expression. Using sva, we employed two surrogate variables with the viral count model. We then used apeglm to shrink the effect sizes and decrease the number of false positive associations. We utilized the EnhancedVolcano package v1.14.0 to represent the resulting data, shown in **Figure 1a**.

#### 2. Individual subject pathway analysis (z-score analysis) methodology

**Sample filtering**: Samples were excluded from analysis if their total human sequence read count was less than 10e06. Number of subjects passing requirements: 486. Subject counts used in this analysis were grouped by viral PCR categories; High: n = 48, Medium: n = 79, Low: n = 35, None: n = 324. **Batch Correction**: The ComBat_seq function in sva package v3.44.0 (https://doi.org/doi:10.18129/B9.bioc.sva) was employed for batch correction versus the plateID (see metadata). **Generation of z-scores**: Gene expression from nasal swabs on individuals with positive results for PCR-detectable SARS-CoV-2 virus (“High”, “Medium”, “Low”) were compared to the group of negative result individuals (“None”). Each gene expression vector from a positive PCR subject was individually transformed into a gene expression z-score vector by individual comparison specifically with the “None” group gene expression vectors. Each “None” subject’s gene expression z-score vector was similarly calculated vs. the rest of the “None” group’s gene expression vectors. Z-scores in the vectors therefore represent each gene’s expression deviation vs. that gene from the “None” group. The resulting gene expression z-score matrix was used to generate single-subject GSEA analyses. **Subject gene set enrichment analyses (GSEA)**: For each subject, the gene expression z-score vector is ranked from the most positive to the most negative z-score. Each subject’s gene expression z-score vector was used as input into the R package fgsea (fast gene set enrichment analysis) v 1.22.0^110^. To measure the relative enrichment of gene sets for each individual subject based on the z-score rankings, we input each subject’s z-score vector into the fgsea package. The MSigDB pathway database version 7.5.1 (doi: https://davislaboratory.github.io/msigdb) was employed as gene sets in the fgsea analysis, specifically the Hallmarks (H) and computational pathways (c2.cp) pathway sets. Results are shown in **Figure 1c** and in supplementary data.

## Acknowledgments

This work was supported by the Division of Intramural Research, NIAID, NIH grant to S.M.B, and DOD W81XWH-21-1-0128 grant awarded to D.C.W. This work was also supported, in whole or in part, by the Bill & Melinda Gates Foundation Grant# INV-046722 awarded to D.C.W. Under the grant conditions of the Foundation, a Creative Commons Attribution 4.0 Generic License has already been assigned to the Author Accepted Manuscript version that might arise from this submission.

This work was also supported by awards from the Defense Health Program (HU00012020067 and HU00012120103), the National Institute of Allergy and Infectious Disease (HU00011920111), and the USU RESPONSE award (HU00012020070). The EPICC protocol was executed by the Infectious Disease Clinical Research Program (IDCRP), a Department of Defense (DoD) program executed by the Uniformed Services University of the Health Sciences (USUHS) through a cooperative agreement by the Henry M. Jackson Foundation for the Advancement of Military Medicine, Inc. (HJF). This project has been funded in part by the National Institute of Allergy and Infectious Diseases at the National Institutes of Health, under an interagency agreement (Y1-AI-5072). We thank the members of the EPICC COVID-19 Cohort Study Group for their many contributions in conducting the study and ensuring effective protocol operations.

## Disclaimer

The opinions expressed in this article are those of the authors and do not reflect the views of the National Institutes of Health, the Department of Health and Human Services, and the United States Government. The contents of this publication are the sole responsibility of the author (s) and do not necessarily reflect the views, opinions, or policies of Uniformed Services University of the Health Sciences (USUHS); the Department of Defense (DoD); the Departments of the Army, Navy, or Air Force; the Defense Health Agency: the Henry M. Jackson Foundation for the Advancement of Military Medicine Inc; the National Institutes of Health. Mention of trade names, commercial products, or organizations does not imply endorsement by the U.S. Government. The investigators have adhered to the policies for the protection of human subjects as prescribed in 45 CFR 46.

## Other Affiliations

R.E.S. is on the scientific advisory of Miromatrix, Inc. and Lime Therapeutics and is a consultant for Alnylam, Inc. D.C.W. is on the scientific advisory boards of Pano Therapeutics, Inc., and Medical Excellent Capital.

## Potential Conflicts of Interest

S.P., M.S., T.B, and D.T. report that the Uniformed Services University (USU) Infectious Diseases Clinical Research Program (IDCRP), a US Department of Defense institution, and the Henry M. Jackson Foundation for the Advancement of Military Medicine, Inc (HJF) were funded under a Cooperative Research and Development Agreement to conduct an unrelated phase III COVID-19 monoclonal antibody immunoprophylaxis trial sponsored by AstraZeneca. The HJF, in support of the USU IDCRP, was funded by the Department of Defense Joint Program Executive Office for Chemical, Biological, Radiological, and Nuclear Defense to augment the conduct of an unrelated phase III vaccine trial sponsored by AstraZeneca. Both trials were part of the US Government’s COVID-19 response. Neither is related to the work presented here. All other authors declare no competing interests.

## Author contributions

S.B.B. and A.B. conceived the project. J.W.G, M.T., A.B., S.B.B., and D.C.W. directed the project. A.B. and D.T. performed the computational analysis with input from J.W.G., K.B., and H.C. P.G. performed the miR-2392 mimic experiments in 3D HUVEC-MT cells. J.W.G., K.B., and J.H. assisted in statistical and computational analysis. J. Frere generated SARS-CoV-2–infected hamster data. C.M., J. Foox, Y.B., R.E.S., and C.E.M. generated the human nasopharyngeal and autopsy tissue transcription data. A.C., A.B., S.S., C.L., and R.E.S. generated the human lymph node histology data. M.T.H., N.J.M., V.K.B., E.A.M., S.A.T.-B., E.J.A., W.A.S., R.J.D., and J.C.S. generated the SARS-CoV-2 MA10–infected C57BL/6 and BALB/c mouse lung transcription data. E.W. and J.H. found and provided pre-processing for the PBMC transcription data, figures/analysis generated by A.B. and input from J.W.G. N.E., B.A., J.C., M.S., D.T., T.B., C.D., and S.P. generated the Whole Blood transcription data. Specific analysis and figure generation on Whole Blood data by A.B. J.W.G., M.T., S.B.B., and D.C.W. wrote the manuscript and A.B. E.S.W., N.S.T., K.B.C, V.Z., P.M.M-V., E.W., S.P., D.T., R.E.S. and D.C.W, participated in regular discussions and helped with editing the manuscript. All authors read and reviewed the final version of the manuscript.

## GROUP AUTHOR BLOCK (PUBMED index)

We thank the members of the EPICC COVID-19 Cohort Study Group for their many contributions in conducting the study and ensuring effective protocol operations. The following members were all closely involved with the design, implementation, and/or oversight of the study and have met group authorship criteria for this manuscript: *Alexander T. Augusta Military Medical Center, VA, USA*: Derek Larson; *Uniformed Services University of the Health Sciences, Bethesda, MD*: Margaret Sanchez, Ed Parmelee, Julia Rozman, Amber Michel, Robert O’Connell, Andrew Snow, Paul Blair; *Madigan Army Medical Center, Joint Base Lewis McChord, WA*: Christopher Colombo, Rhonda Colombo; *Naval Medical Center Portsmouth, Portsmouth, VA, USA*: Tahaniyat Lalani; *Naval Medical Center San Diego, San Diego, CA, USA:* Ryan Maves, Catherine Berjohn; *Tripler Army Medical Center, Honolulu, HI, USA:* Milissa Jones; *U.S. Air Force School of Aerospace Medicine, Wright-Patterson, OH, USA:* Anthony Fries; *Walter Reed National Military Medical Center, Bethesda, MD:* Anuradha Ganesan; *William Beaumont Army Medical Center, El Paso, TX, USA*: Rupal Mody

## Data and material availability

The published article includes all datasets generated and analyzed during this study. Processed bulk RNA-seq data for the human-related data from the nasopharyngeal and autopsy data is available online with dbGaP Study Accession number: phs002258.v1.p1 and online here at:https://www.ncbi.nlm.nih.gov/projects/gap/cgi-bin/study.cgi?study_id=phs002258.v1.p1 and also https://covidgenes.weill.cornell.edu/. The murine RNA-seq data is deposited on the Gene Expression Omnibus (GEO) repository with accession number GSE221510 and can be found here: https://www.ncbi.nlm.nih.gov/geo/query/acc.cgi?acc=GSE221510. Additionally, the murine RNA-seq data was also deposited in the NCBI BioProject database (https://www.ncbi.nlm.nih.gov/bioproject/) under the BioProject accession number PRJNA803057. The hamster RNA-seq data is also deposited on the GEO repository with accession number GSE203001 and can be found here: https://www.ncbi.nlm.nih.gov/geo/query/acc.cgi?acc=GSE203001. The RNA-seq data from the miR-2392 mimic 3D HUVEC tissue model data is available via the NASA Open Science Data Repository’s (OSDR) Biological Data Management Environment (https://osdr.nasa.gov/bio/repo) with accession number: OSD-577 and DOI: 10.26030/rs3g-e189. This study did not generate new unique reagents.

## SUPPLEMENTAL MATERIALS

Figures S1 to S10

Tables S1 to S7

## Notes

### Summary of Updates

Authors and affiliations were updated.

