## Supplemental Data for "Lethal COVID-19 Associates With RAAS-Induced Inflammation For Multiple Organ Damage Including Mediastinal Lymph Nodes"

**Figure S1.**

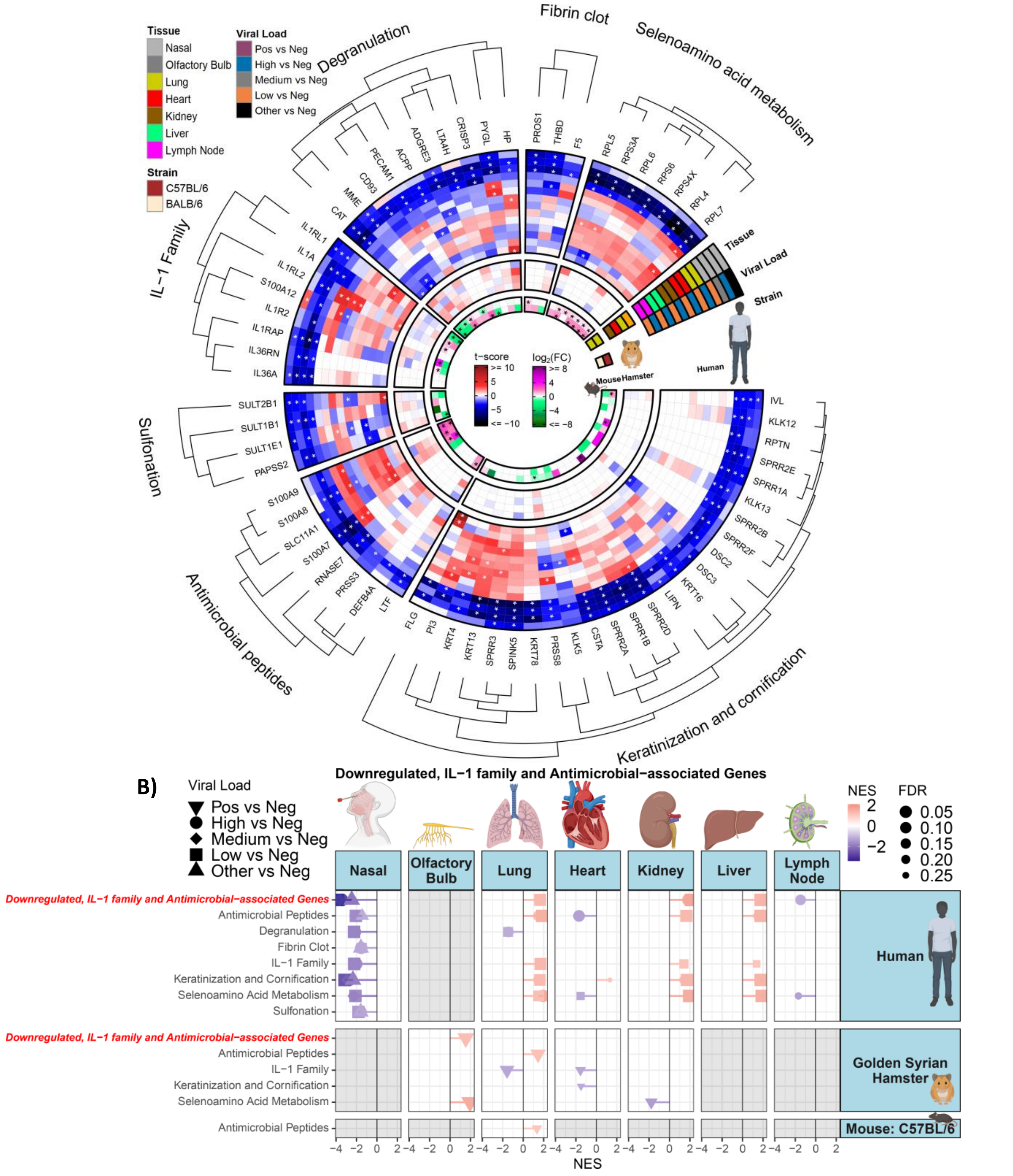


**Fig. S1: Differentially regulated downregulated genes of interest in infected rodent samples and human autopsy tissues from COVID-19 patients. A)** Circular heatmap displaying the t-score statistics for specific downregulated genes of interest, comparing SARS-CoV-2 infected hamster tissues, viral load versus negative patient nasopharyngeal and autopsy tissue samples, and $log_2$-foldchange (FC) for SARS-CoV-2 infected C57BL/6 and BALB/c mouse lungs. **B)** Lollipop plots for statistically significant changes in downregulated genes of interest sets determined by fGSEA for SARS-CoV-2 positive nasopharyngeal, autopsy, and rodent tissues, ranked by NES, nominal enrichment score.

**Figure S2.**

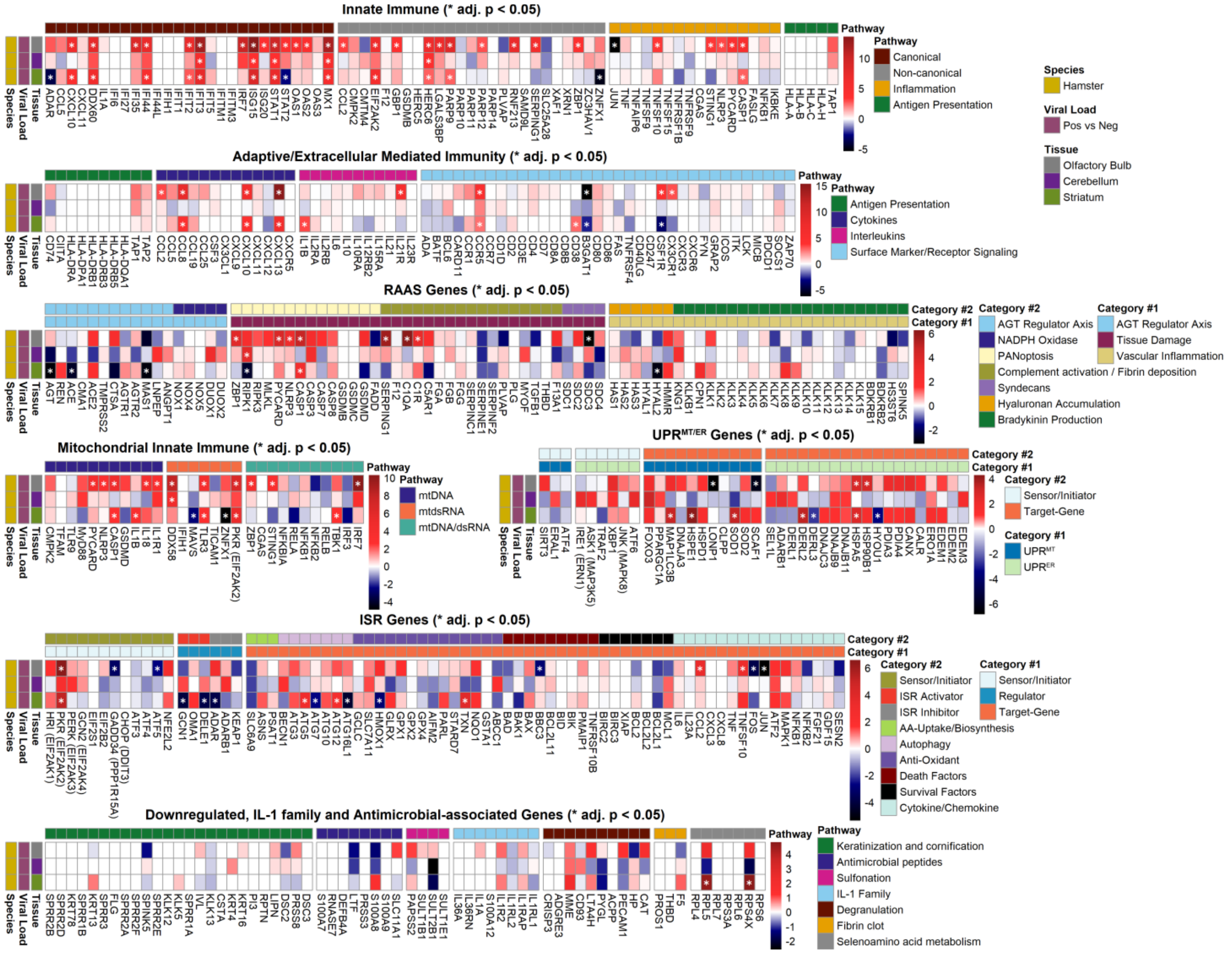

**Fig. S2: Differentially regulated immune genes in SARS-CoV-2 infected hamster olfactory bulb, cerebellum, and striatum.** Linear heatmap displaying the t-score statistics for innate immune genes of interest, comparing SARS-CoV-2 infected hamster tissues

**Figure S3.**

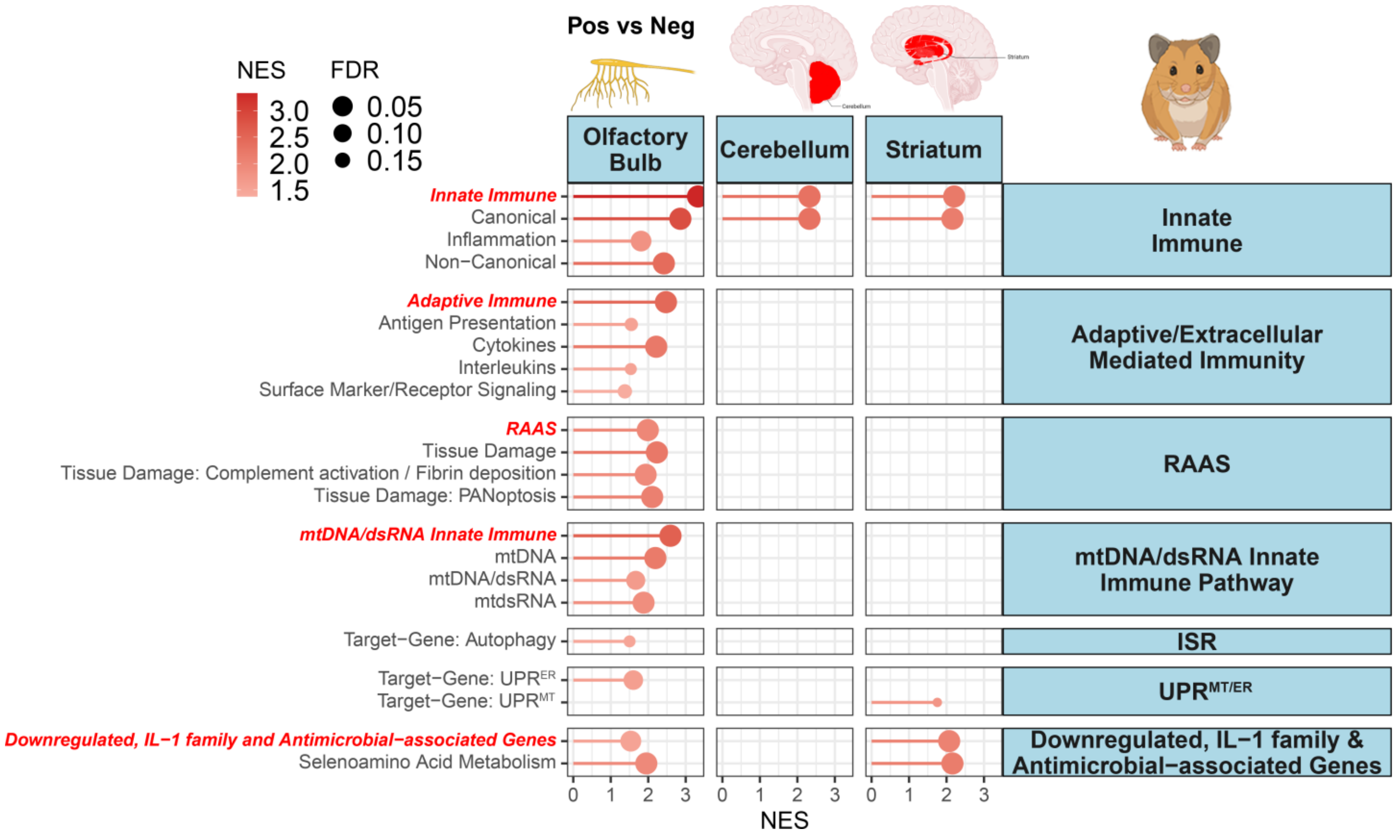


**Fig. S3: Lollipop plots for statistically significant changes in innate immune gene sets determined by fGSEA for SARS-CoV-2 positive hamster tissues**. Ranked by NES, nominal enrichment score.

**Figure S4.**

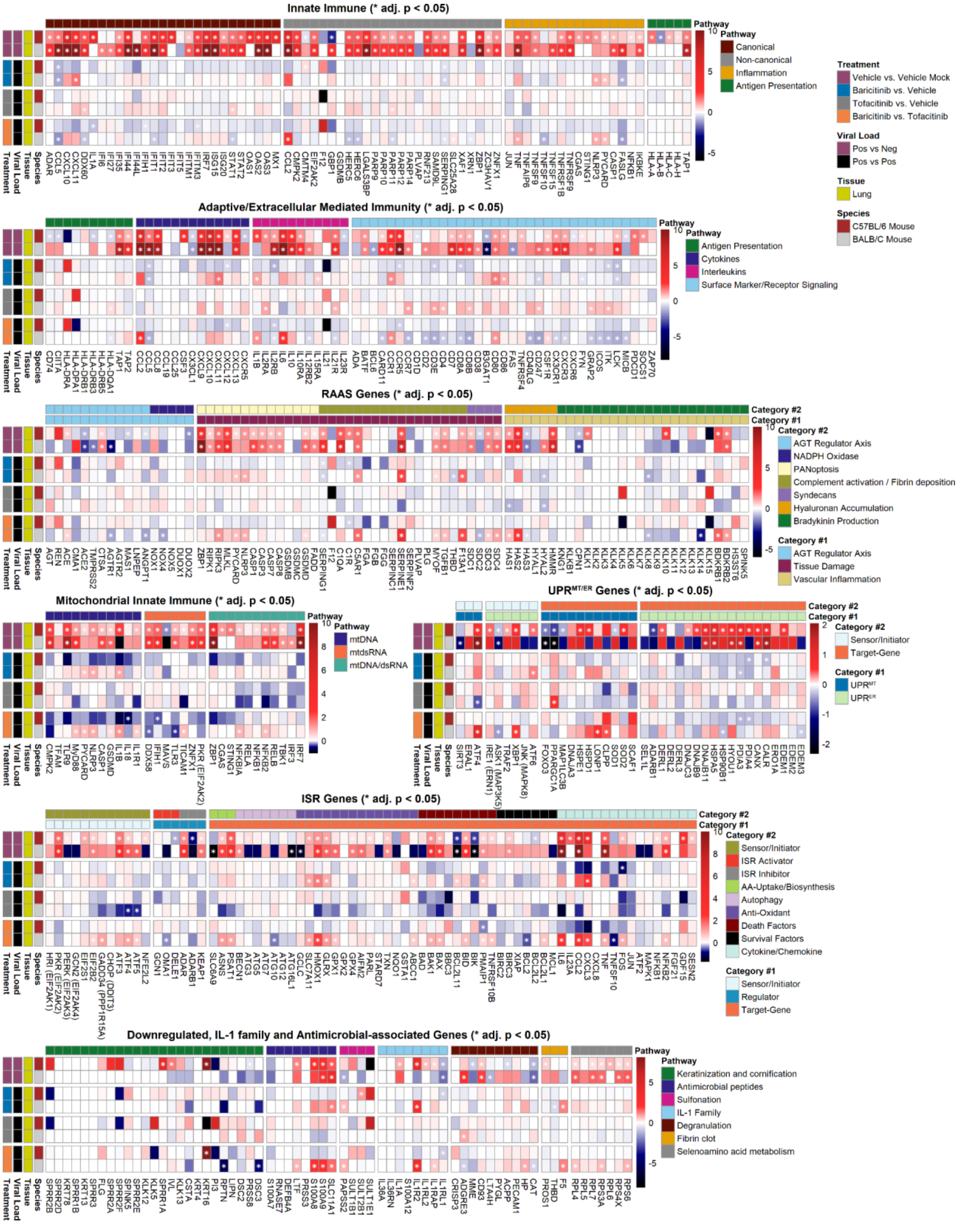


**Fig. S4: Differentially regulated immune genes in SARS-CoV-2 infected C57BL/6 and BALB/c mouse lungs treated with or without baricitinib or tofacitinib.** Linear heatmap displaying the t-score statistics for innate immune genes of interest, comparing SARS-CoV-2 infected mouse lungs treated with or without baricitinib or tofacitinib.

**Figure S5.**

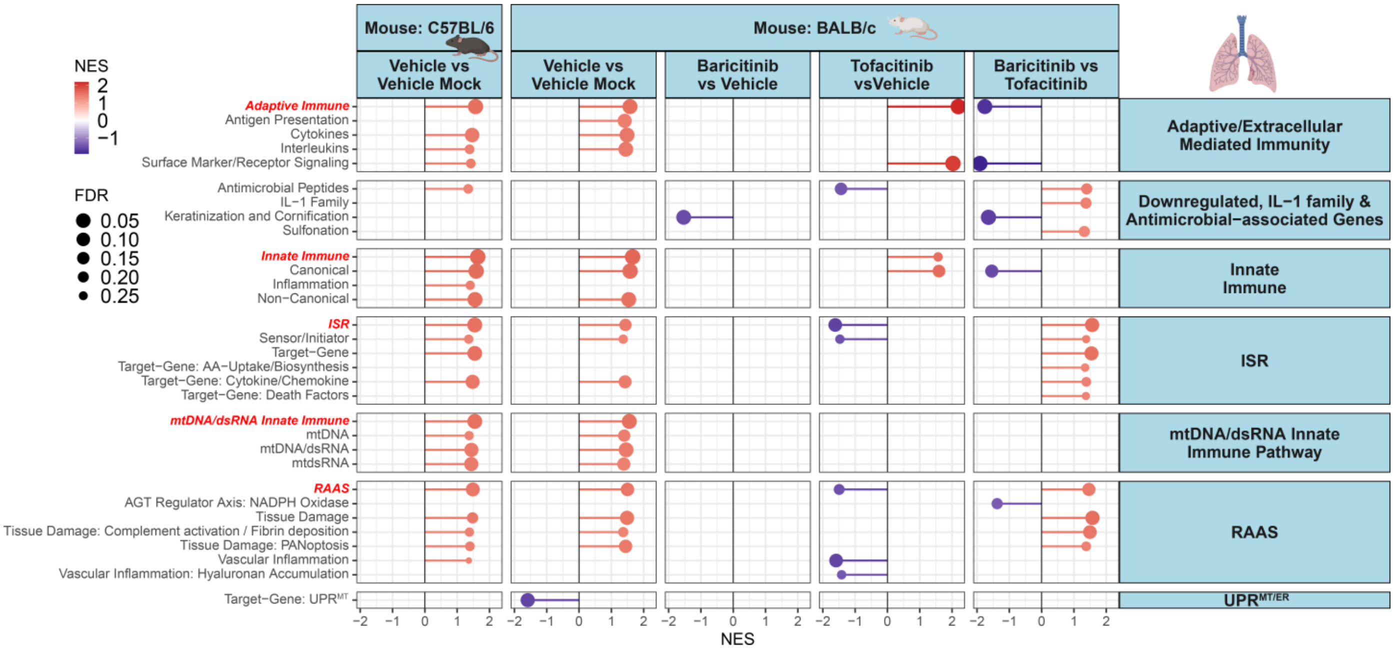


**Fig. S5: Lollipop plots for statistically significant changes in innate immune gene sets determined by fGSEA for SARS-CoV-2 positive mouse lungs treated with or without baricitinib or tofacitinib.** Ranked by NES, nominal enrichment score.

**Figure S6.**

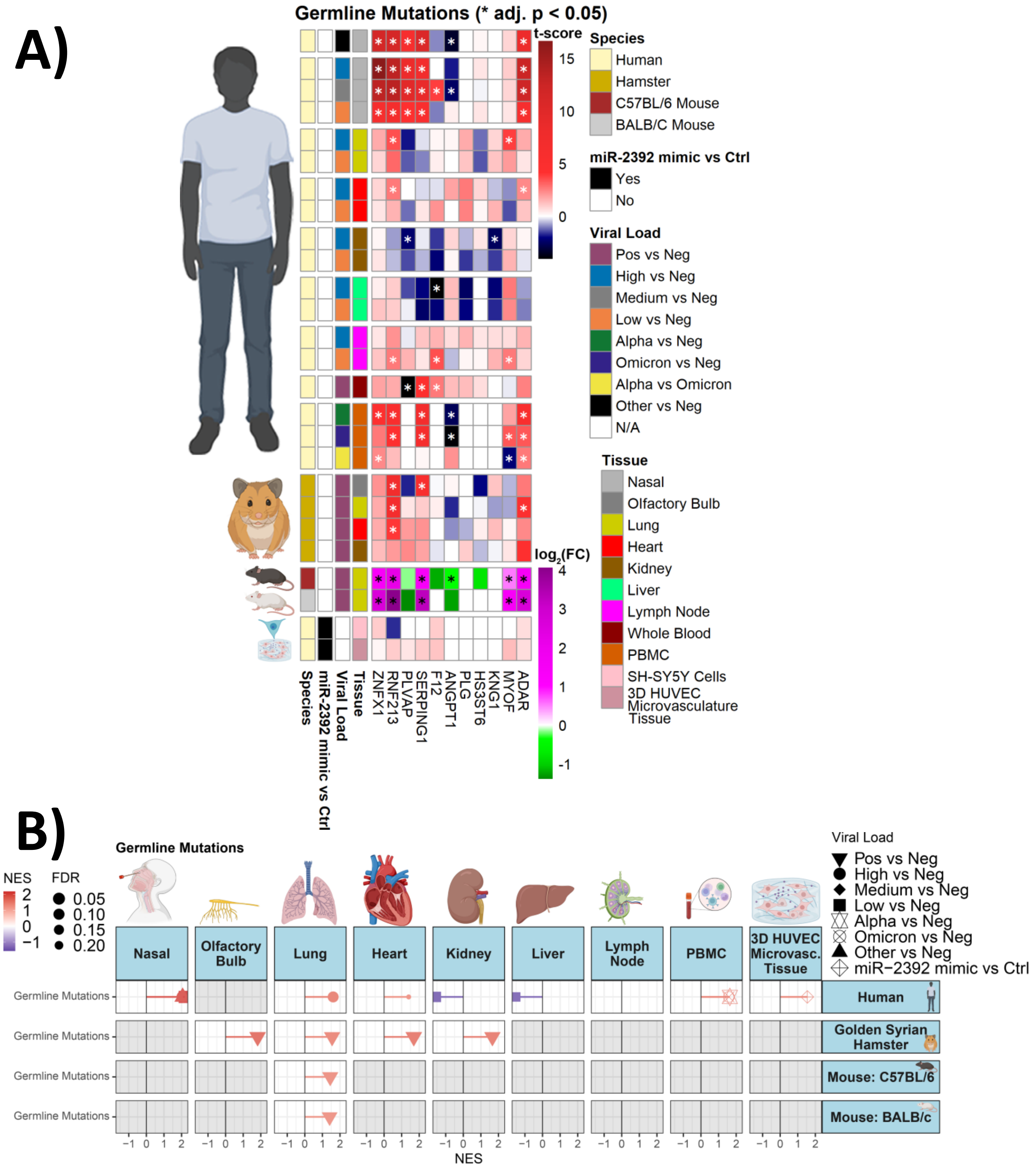


**Fig. S6: Differentially regulated genes that have germline mutations in infected rodent samples and human autopsy and nasopharyngeal tissues from COVID-19 patients.** Linear heatmap displaying the t-score statistics for genes that have germline mutations of interest, comparing SARS-CoV-2 infected hamster tissues, viral load versus negative patient nasopharyngeal and autopsy tissue samples, and $log_2$-foldchange (FC) for SARS-CoV-2 infected C57BL/6 and BALB/c mouse lungs. **B)** Lollipop plots for statistically significant changes in germline mutation gene sets determined by fGSEA for SARS-CoV-2 positive nasopharyngeal, autopsy, and rodent tissues, ranked by NES, nominal enrichment score.

**Figure S7.**

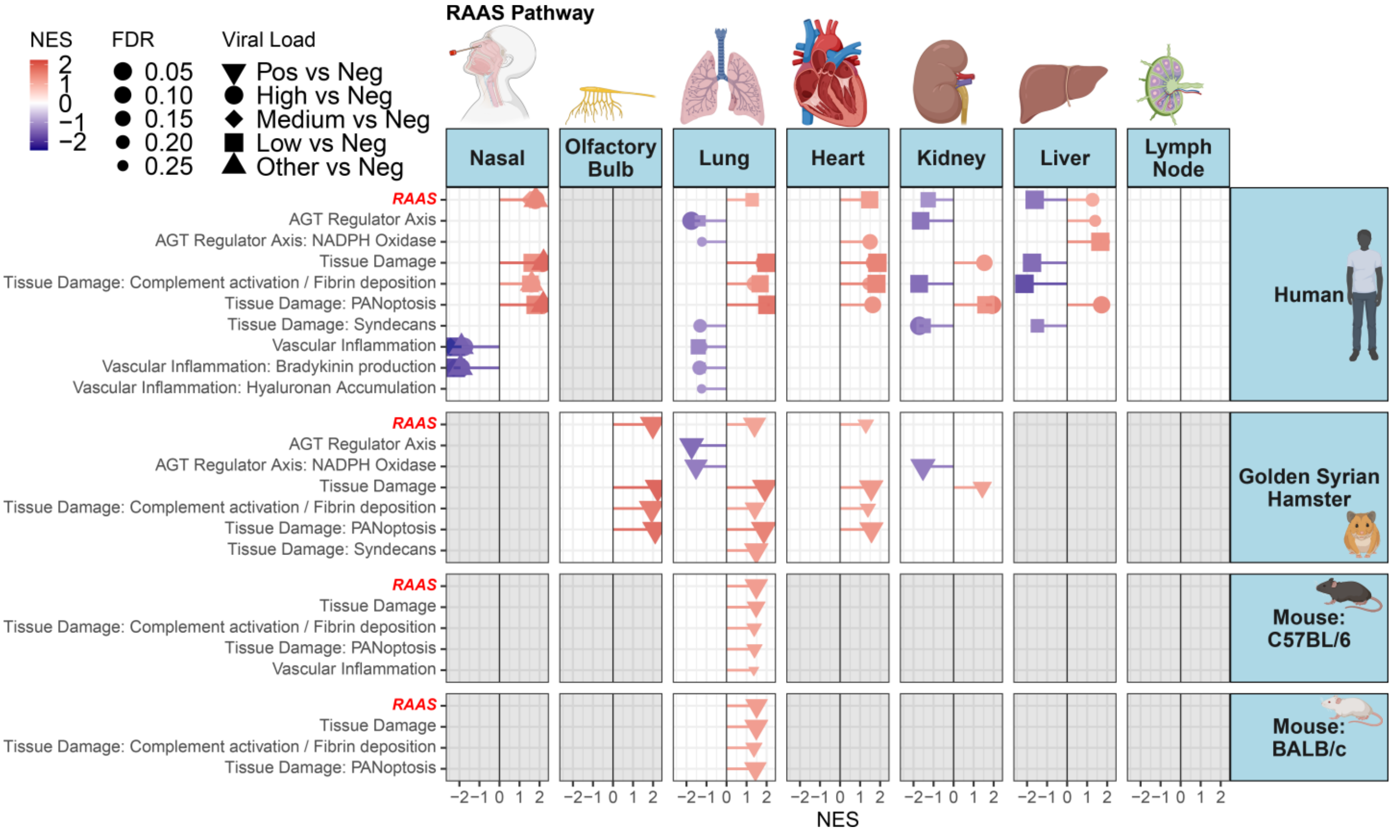


**Fig. S7: Lollipop plots for statistically significant changes in RAAS gene sets determined by fGSEA in SARS-CoV-2 infected rodent samples and human autopsy and nasopharyngeal tissues from COVID-19 patients.** Ranked by NES, nominal enrichment score.

**Figure S8.**

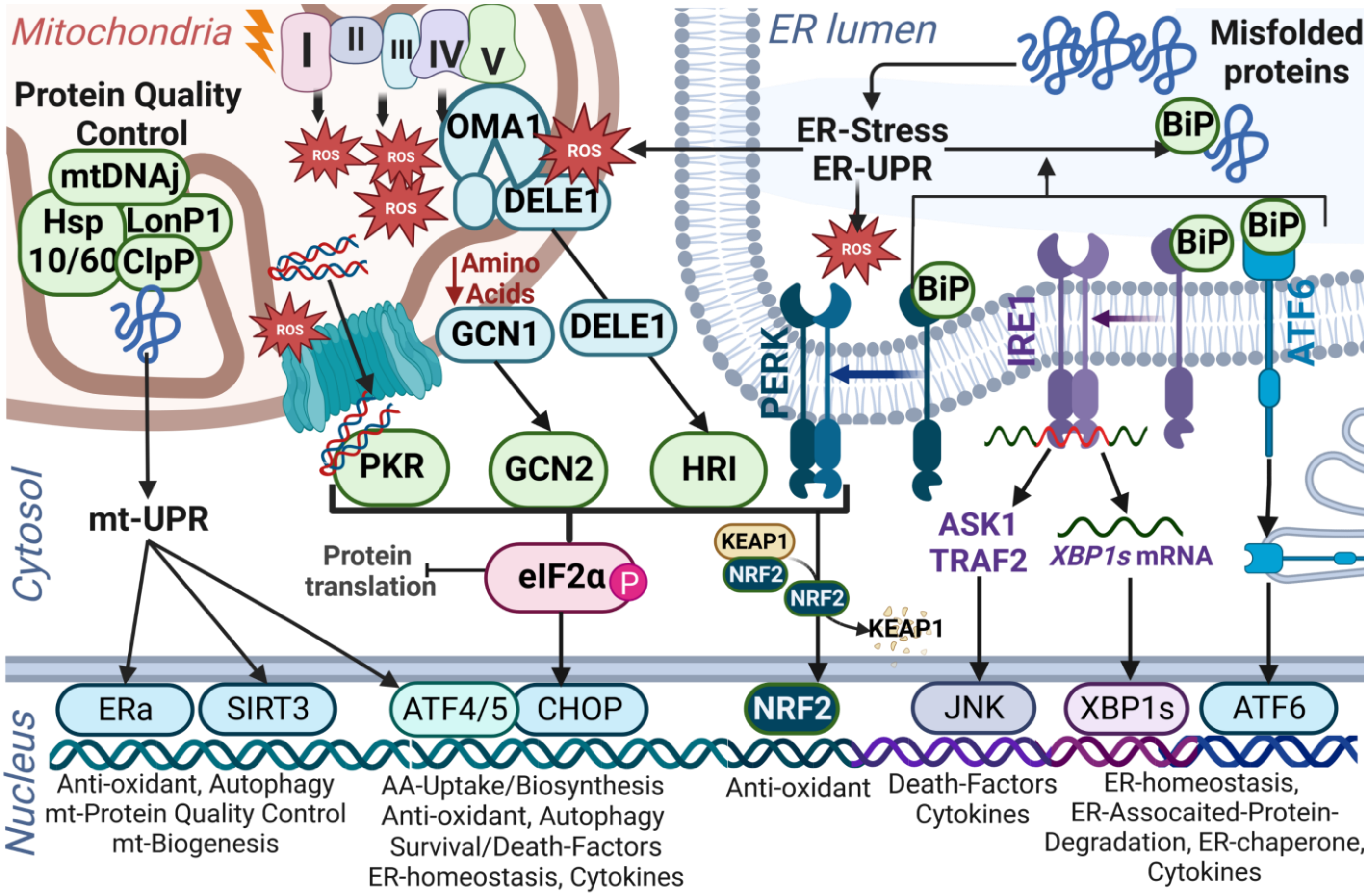


**Fig. S8: Summary of key ISR, endoplasmic reticulum and mitochondrial UPR genes.**

**Figure S9.**

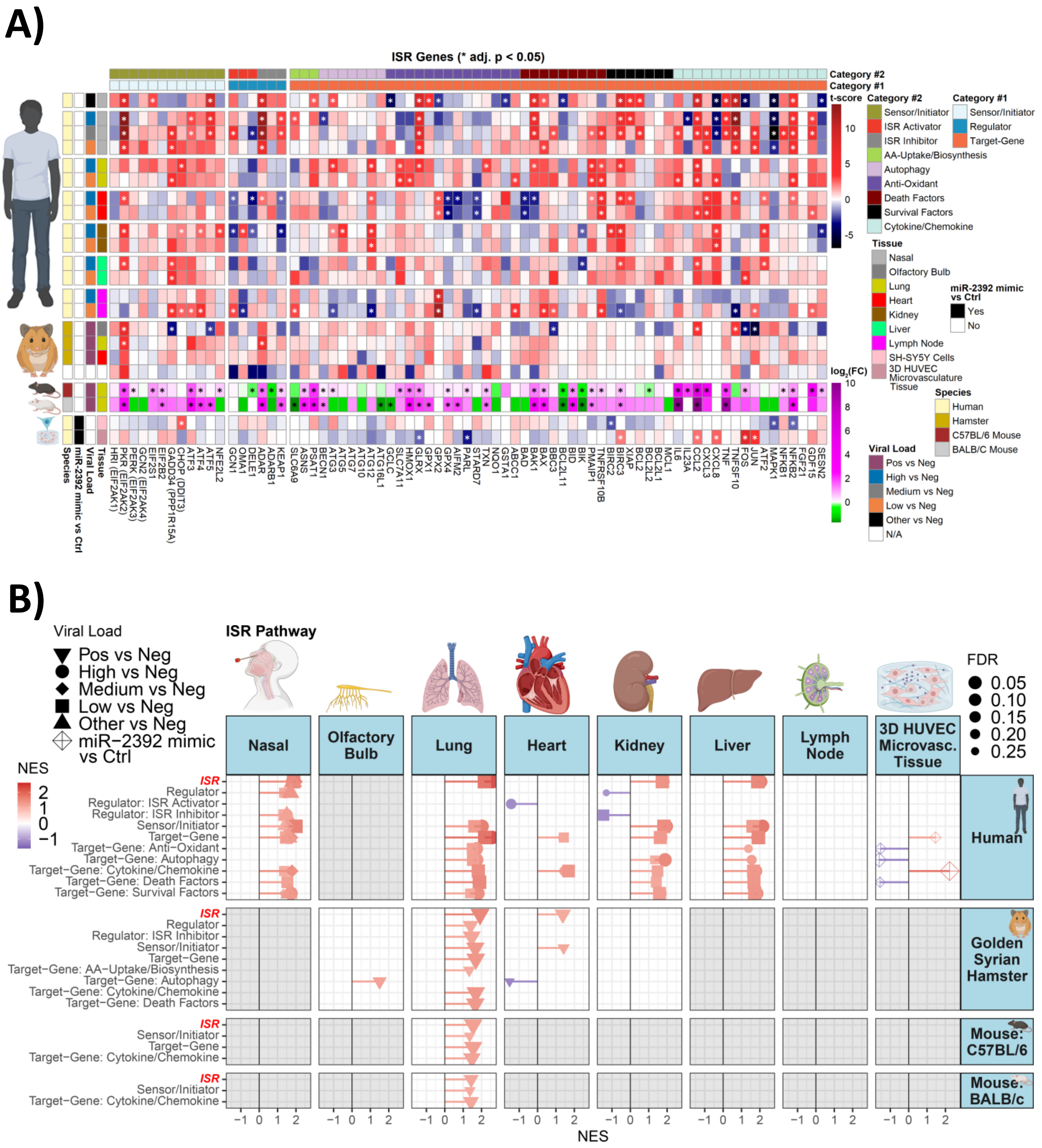


**Fig. S9: Differentially regulated ISR genes in SARS-CoV-2 infected rodent samples and human autopsy and nasopharyngeal tissues from COVID-19 patients. A)** Linear heatmap displaying the t-score statistics specific ISR gene sets, comparing SARS-CoV-2 infected hamster tissues, viral load versus negative patient nasopharyngeal and autopsy tissue samples, and $log_2$-foldchange (FC) for SARS-CoV-2 infected C57BL/6 and BALB/c mouse lungs. **B)** Lollipop plots for statistically significant changes in custom ISR gene sets determined by fGSEA for SARS-CoV-2 positive nasopharyngeal, autopsy, and rodent tissues, and miR-2392-expressing 3D-HUVEC-MT cells ranked by NES, nominal enrichment score.

**Figure S10.**

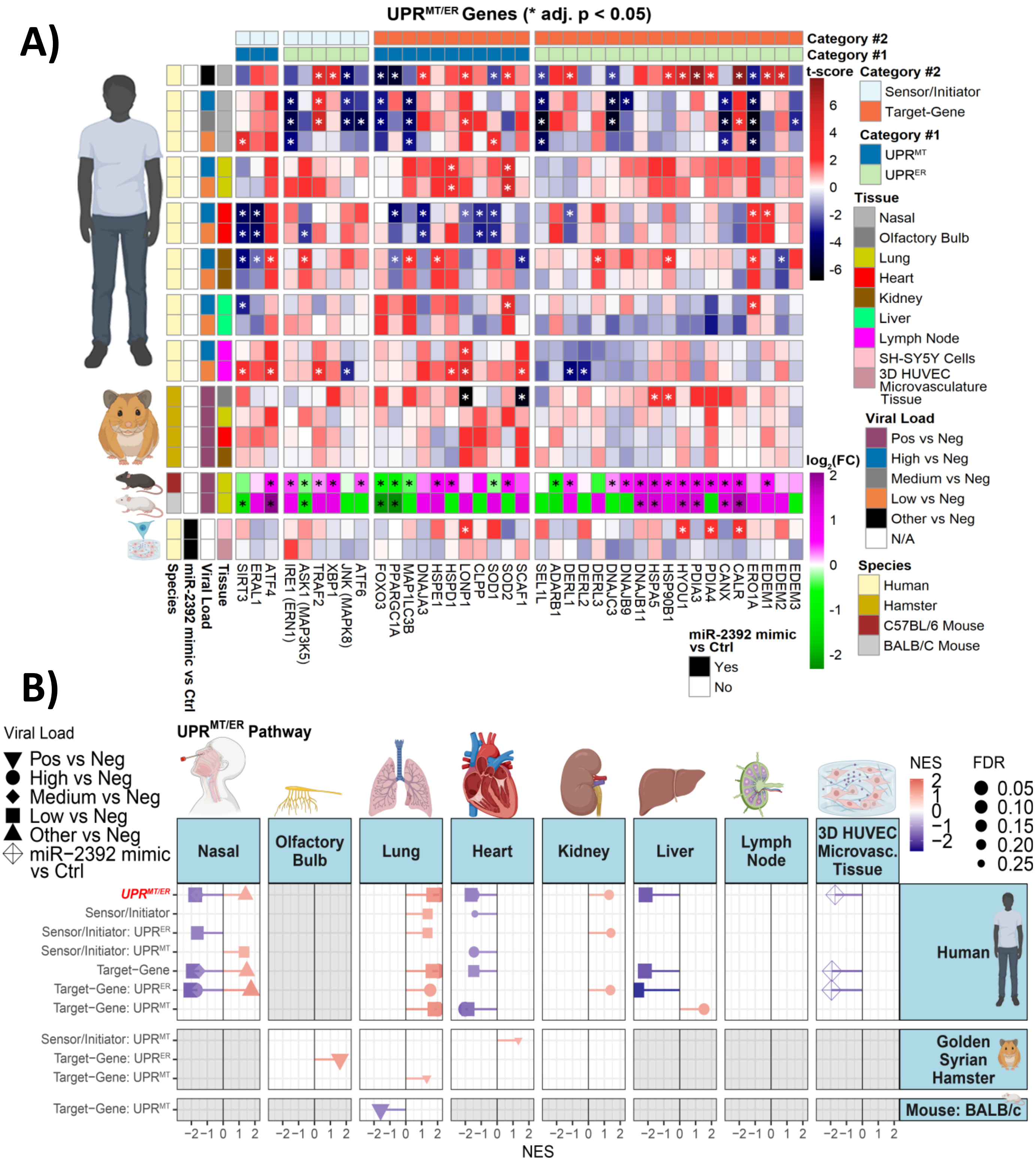


**Fig. S10: Differentially regulated UPR$^{ER/MT}$ genes in SARS-CoV-2 infected rodent samples and human autopsy and nasopharyngeal tissues from COVID-19 patients. A)** Linear heatmap displaying the t-score statistics specific UPR$^{ER/MT}$ gene sets, comparing SARS-CoV-2 infected hamster tissues, viral load versus negative patient nasopharyngeal and autopsy tissue samples, and $log_2$-foldchange (FC) for SARS-CoV-2 infected C57BL/6 and BALB/c mouse lungs. **B)** Lollipop plots for statistically significant changes in custom UPR$^{ER/MT}$ gene sets determined by fGSEA for SARS-CoV-2 positive nasopharyngeal, autopsy, and rodent tissues, and miR-2392-expressing 3D-HUVEC-MT cells ranked by NES, nominal enrichment score.

**Figure S11.**

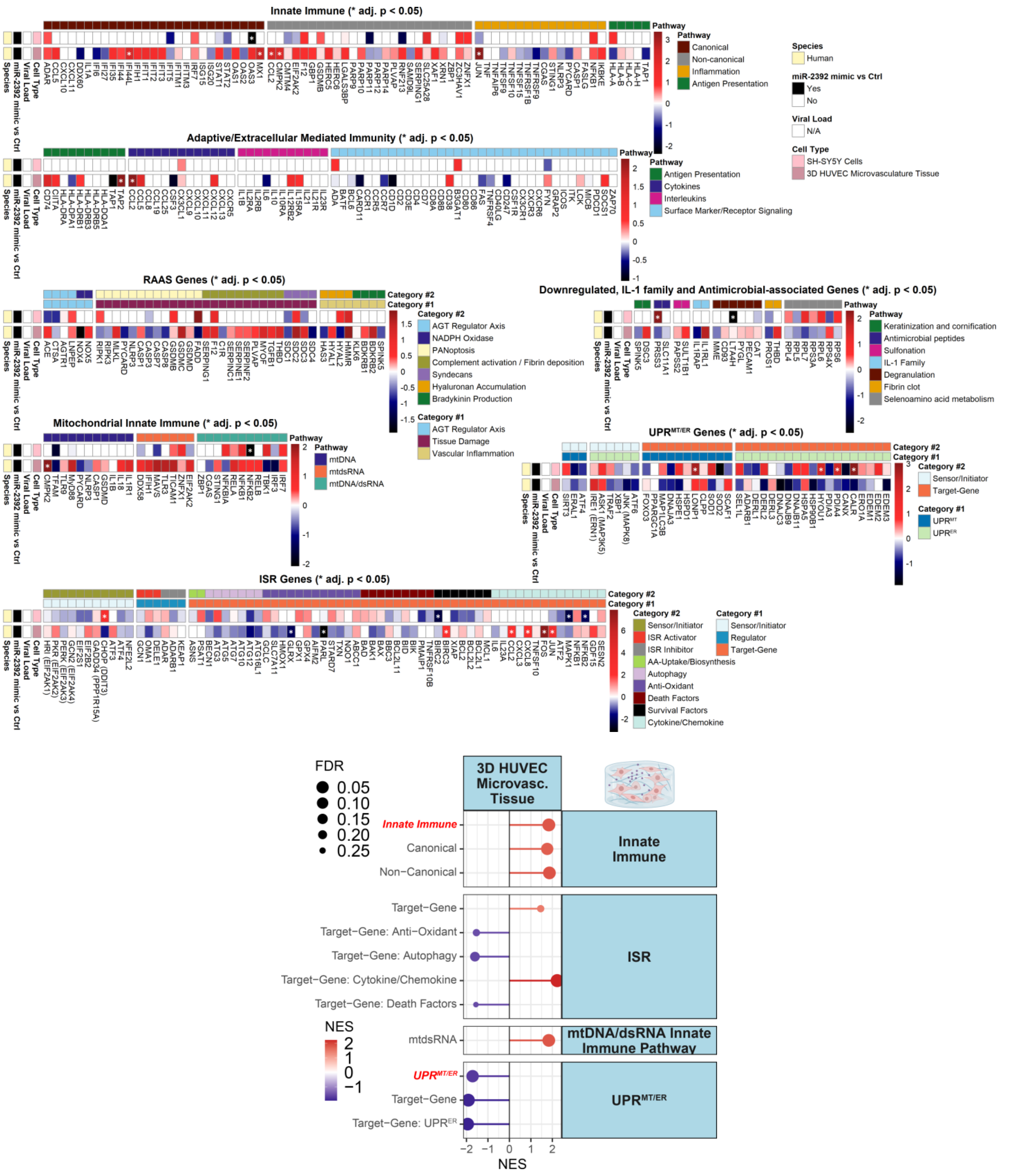


**Fig. S11: Differentially regulated immune genes in miR-2392-expressing 3D-HUVEC-MT cells. A)** Linear heatmap displaying the t-score statistics for innate immune genes comparing miR-2392-expressing 3D-HUVEC-MT cells. **B)** Lollipop plots for statistically significant changes in innate immune gene sets determined by miR-2392-expressing 3D-HUVEC-MT cells, ranked by NES, nominal enrichment score.

**Figure S12.**

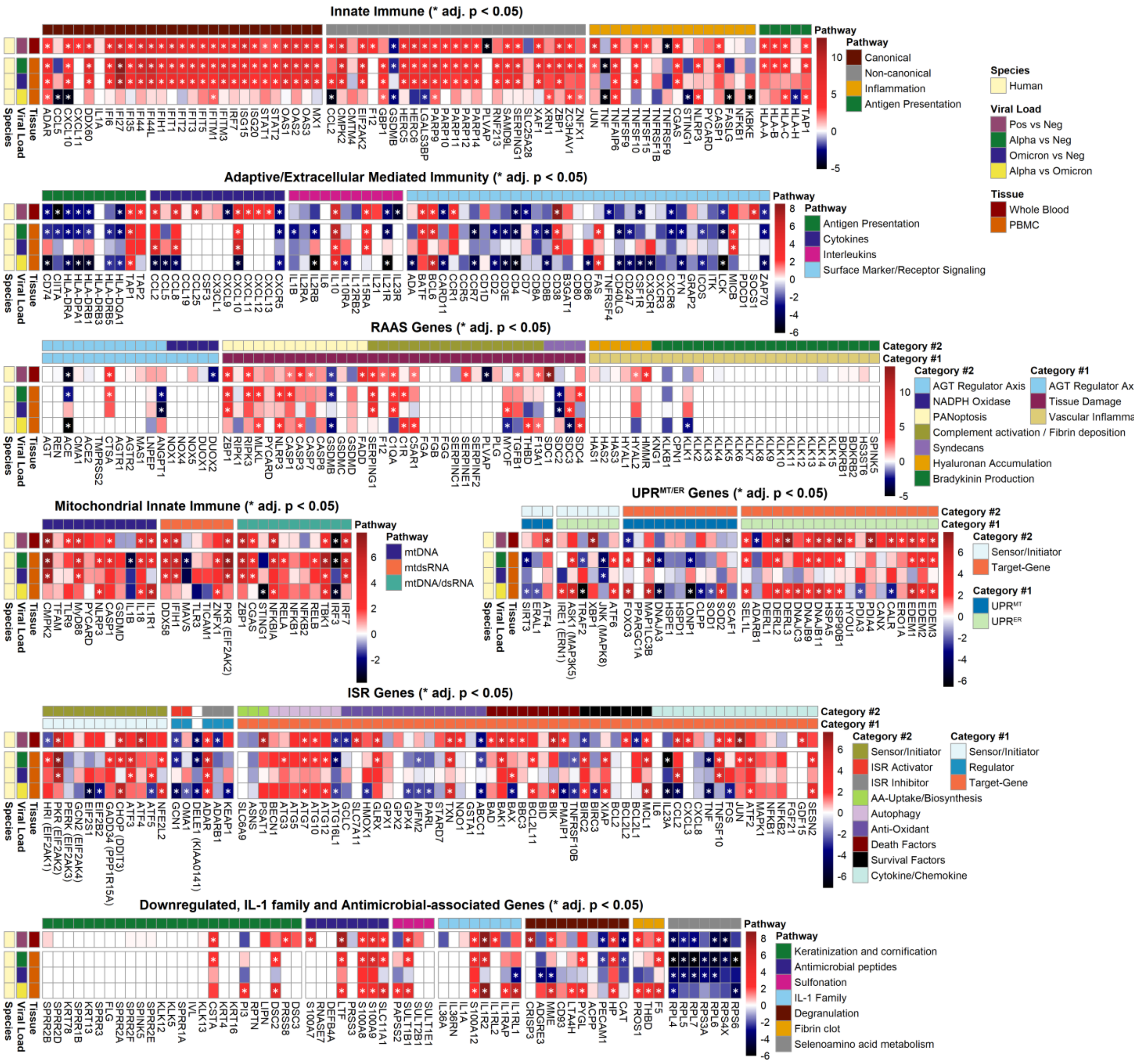


**Fig. S12: Differentially regulated immune genes in PBMCs and whole blood samples collected from SARS-CoV-2-Alpha and SARS-CoV-2-Omicron-infected patients.** Linear heatmap displaying the t-score statistics for innate immune genes comparing PBMCs and whole blood samples collected from SARS-CoV-2-Alpha and SARS-CoV-2-Omicron-infected patients.

**Table. S1. SARS-CoV-2 Viral Load of Human Nasopharyngeal, Autopsy, Rodent and PBMCs Samples.**

| STAGE | Sample | Value | PMID |
|---|---|---|---|
| ACUTE | **Hamster Lungs PFU Log10 ($\bar{X}\pm\sigma_M$)** | | 35857629 |
| | DAY 1 | 7.4 ± 0.2 | |
| | DAY 2 | 8.2 ± 0.4 | |
| | DAY 3 | 7.2 ± 0.3 | |
| | DAY 7 | 0 ± 0 | |
| | **Hamster Tissues 3DPI TPM ($\bar{X}\pm\sigma_M$)** | | 35233572 |
| | Lung | 2.3E+05 ± 4.0E+04 | |
| | Kidney | 8 ± 8 | |
| | Heart | 58 ± 16 | |
| | Olfactory Bulb | 581 ± 247 | |
| | Cerebellum | 428 ± 220 | |
| | Striatum | 75 ± 73 | |
| EARLY | **Nasal Samples TPM ($\bar{X}\pm\sigma_M$)** | | |
| | Nasal-High | 1.0E+07 ± 1.4E+06 | |
| | Nasal-Medium | 7.0E+05 ± 1.6E+05 | |
| | Nasal-Low | 1.5E+04 ± 1.0E+04 | |
| | Other-Viral | 18 ± 5 | |
| MID | **Mouse Lungs 4DPI PFU Log10 ($\bar{X}\pm\sigma_M$)** | | |
| | C57BL/6 | 4.76 ± 4.8 | |
| | BALB/c | 4.85 ± 4.6 | |
| LATE | **Autopsy Tissue TPM ($\bar{X}\pm\sigma_M$)** | | |
| | Lung | 1 ± 1 | |
| | Heart | 1 ± 1 | |
| | Kidney | 1 ± 1 | |
| | Liver | 1 ± 1 | |
| | LymphNode | 2 ± 2 | |
| | **PBMCs TPM ($\bar{X}\pm\sigma_M$)** | | Haltman, J (2023), in press |
| | PBMCs α-strain | 0 ± 0 | |
| | PBMCs o-strain | 0 ± 0 | |

**Table S2. Curated Immune Gene List**

### INNATE IMMUNE GENES

***Canonical:*** ADAR, CCL5, CXCL10, CXCL11, DDX60, IFI27, IFI35, IFI44, IFI44L, IFI6, IFIH1, IFIT1, IFIT2, FIT3, IFIT5, IFITM1, IFITM3, IL1A, IRF7, ISG15, ISG20, MX1, OAS1, OAS2, OAS3, PARP9, SAMD9L, STAT1, STAT2

***Non-Canonical:*** CCL2, CMPK2, CMTM4, EIF2AK2, GBP1, GSDMB, HERC5, HERC6, LGALS3BP, PARP10, PARP11, PARP12, PARP14, PARP9, PLVAP, RNF213, SERPING1, SLC25A28, XAF1, ZBP1, ZC3HAV1, ZNFX1

***Inflammation***: CASP1, CGAS, FASLG, IKBKE, JUN, NFKB1, NLRP3, PYCARD, STING1, TNF, TNFAIP6, TNFRSF1B, TNFRSF9, TNFSF10, TNFSF15, TNFSF9

***Antigen Presentation:*** HLA-A, HLA-B, HLA-C, HLA-H, TAP1

### DOWNREGULATED GENES OF INTEREST

***Keratinization and Cornification:*** CSTA, DSC2, DSC3, FLG, IVL, KLK12, KLK13, KLK5, KRT13, KRT16, KRT4, KRT78, LIPN, PI3, PRSS8, RPTN, SPINK5, SPRR1A, SPRR1B, SPRR2A, SPRR2B, SPRR2D, SPRR2E, SPRR2F, SPRR3

***Antimicrobial Peptides:*** DEFB4A, LTF, PRSS3, RNASE7, S100A7, S100A8, S100A9, SLC11A1

***Sulfonation:*** PAPSS2, SULT1B1, SULT1E1, SULT2B1

***IL-1 Family:*** IL1A, IL1R2, IL1RAP, IL1RL1, IL1RL2, IL36A, IL36RN, S100A12

***Degranulation:*** ACPP, ADGRE3, CAT, CD93, CRISP3, HP, LTA4H, MME, PECAM1, PYGL

***Fibrin Clot:*** F5, PROS1, THBD

***Selenoamino acid Metabolism:*** RPL4, RPL5, RPL6, RPL7, RPS3A, RPS4X, RPS6

### MITOCHONDRIAL INNATE IMMUNE

***mtDNA/dsRNA activated genes:***
***mtDNA:*** CASP1, CMPK2, GSDMD, IL18, IL1B, IL1R1, MYD88, NLRP3, PYCARD, TFAM, TLR9
***mtdsRNA:*** DDX58, EIF2AK2, IFIH1, MAVS, TICAM1, TLR3, ZNFX1
***mtDNA/dsRNA:*** CGAS, IRF3, IRF7, NFKB1, NFKB2, NFKBIA, RELA, RELB, STING1, TBK1, ZBP1

***Integrated Stress Response (ISR):***
***Sensor/Initiator:*** ATF3, ATF4, ATF5, CHOP (DDIT3), EIF2B2, EIF2S1, GADD34 (PPP1R15A), GCN2 (EIF2AK4), HRI (EIF2AK1), NFE2L2, PERK (EIF2AK3), PKR (EIF2AK2)

***Regulator:***
*ISR Activator:* DELE1, GCN1, OMA1
*ISR Inhibitor:* ADAR, ADARB1, KEAP1

***Target-Gene:***
*AA-Uptake/Biosynthesis*: ASNS, SLC6A9, PSAT1
*Autophagy*: BECN1, ATG10, ATG12, ATG3, ATG5, ATG7
*Anti-Oxidant*: ABCC1, AIFM2, GCLC, GCLC, GLRX, GPX1, GPX2, GPX4, GSTA1, HMOX1, NQO1, PARL, SLC7A11, STARD7, TXN
*Death Factors*: BAD, BAK1, BAX, BBC3, BCL2L11, BID, BIK, PMAIP1, TNFRSF10B
*Survival Factors*: BCL2, BCL2L1, BCL2L2, BIRC2, BIRC3, MCL1, XIAP
*Cytokines/Chemokines*: ATF2, CCL2, CXCL3, CXCL8, FGF21, FOS, GDF15, IL23A, IL6, JUN, MAPK1, NFKB1, NFKB2, SESN2, TNF, TNFSF10

***Unfolded Protein Response (UPR):***

***<u>Endoplasmic Reticulum UPR (UPR$^{ER}$):</u>***
<u>Sensor/Initiator</u>: ASK1 (MAP3K5), ATF6, IRE1 (ERN1), JNK (MAPK8), TRAF2, XBP1
<u>Target-Gene</u>: ASK1 (MAP3K5), ATF6, IRE1 (ERN1), JNK (MAPK8), TRAF2, XBP1

***<u>Mitochondrial UPR (UPR$^{MT}$):</u>***
<u>Sensor/Initiator</u>: ERAL1, SIRT3
<u>Target-Gene</u>: CLPP, DNAJA3, FOXO3, HSPD1, HSPE1, LONP1, MAP1LC3B, PPARGC1A, SCAF1, SOD1, SOD2

### EXTRACELLULAR IMMUNITY GENES

***Antigen presentation:*** CD74, CIITA, HLA-DPA1, HLA-DQA1, HLA-DRA, HLA-DRB1, HLA-DRB3, HLA-DRB5, TAP1, TAP2

***Cytokines:*** CCL19, CCL2, CCL25, CCL5, CCL8, CSF3, CX3CL1, CXCL10, CXCL11, CXCL12, CXCL13, CXCL9

***Interleukins:*** IL10, IL10RA, IL12RB2, IL15RA, IL1B, IL21R, IL23R, IL2RA, IL2RB, IL6

***Surface marker/receptor signaling:*** ADA, CARD11, CCR1, CCR5, CCR7, CD1D, CD2, CD247, CD38, CD3E, CD4, CD7, CD80, CD86, CD8A, CD8B, CSF1R, CX3CR1, CXCR3, CXCR6, FYN, GRAP2, ICOS, ITK, LCK, MICB, PDCD1, SOCS1, ZAP70

### RENIN-ANGIOTENSIN-ALDOSTERONE SYSTEM (RAAS)

***AGT Regulator Axis*****:** ACE, ACE2, AGT, AGTR1, AGTR2, ANGPT1, CMA1, CPA3, CTSA, MAS1, REN, TMPRSS2
***<u>NADPH Oxidase</u>*****<u>:</u>** DUOX1, DUOX2, NOX1, NOX4, NOX5

***Tissue Damage:***
***<u>PANoptosis</u>*****<u>:</u>** CASP3, CASP7, CASP8, FADD, GSDMB, GSDMC, GSDMD, MLKL, RIPK1, RIPK3, ZBP1
***<u>Complement activation/Fibrin deposition</u>*****<u>:</u>** C1QA, C1R, C5AR1, F12, F13A1, FGA, FGB, FGG, MYOF, PLG, PLVAP, SERPINC1, SERPINE1, SERPINF2, SERPING1, TGFB1, THBD

***Vascular Inflammation:***
***<u>Hyaluronan Accumulation</u>*****<u>:</u>** HAS1, HAS2, HAS3, HMMR, HYAL1, HYAL2
***<u>Bradykinin production</u>*****<u>:</u>** BDKRB1, BDKRB2, CPN1, HS3ST6, KLK1, KLK2, KLK3, KLK4, KLK5, KLK6, KLK7, KLK8, KLK9, KLK10, KLK11, KLK12, KLK13, KLK14, KLK15, KLKB1, KNG1

### Table S3. Special Mention of Germline Mutations

| Gene Symbol | Germline Mutation | Disease | Description/Function | PMID |
|---|---|---|---|---|
| **ZNFX1** | Reduced Expression Mutations | Aicardi–Goutières syndrome (AGS), when exposed to pathogens | Zinc Finger NFX1-Type Containing 1 (ZNFX1), encodes a zinc finger helicase, ISG protein that recognizes viral dsRNA and shuttles to the outer mitochondrial membrane. ZNFX1 when exposed to pathogens, leads to constitutive IFN-signaling with lethal effects | **34708404** |
| **RNF213** | c.14429G > A (p.Arg4810Lys) Polymorphism | MOYA-MOYA disease | Ring Finger Protein 213 (RNF213), encodes a C3HC4-type RING finger domain that can inhibit the Wnt signaling pathway triggering vessel regression, facilitates Class I MHC-mediated antigen processing and presentation, and is associated with intracranial artery disorders and MOYA-MOYA disease. Symptoms of MOYA-MOYA disease include Headache, seizures, weakness, numbness or paralysis in the face, arm or leg typically on one side of your body, vision problems, trouble speaking or understanding others, known as aphasia, cognitive or developmental delays, involuntary movements. | **32686731** |
| **PLVAP** | Reduced Expression Mutations | Protein losing enterophapy (PLE) | Plasmalemma Vesicle Associated Protein (PLVAP), encodes for an endothelial cell-specific membrane protein required to maintain the integrity of diaphragm structures that bridge endothelial fenestrae, vital for controlling microvascular permeability and barrier function of gut endothelium; its loss in mice can trigger major leaks of fluid and proteins into surrounding tissues, resulting in premature death caused by severe diarrhea, and concomitant edema of the intestine, kidneys, and pancreas. Reduced expression mutations in PLVAP, results in protein losing enterophapy (PLE) in the gut. Symptoms of PLE include diarrhea, tissue swelling (edema), ascites (excess fluid trapped in your abdomen), pleural and pericardial effusions (excess fluid around your heart), hypoproteinemia (lower than normal protein levels in your body), severe malnutrition. | **36781482. 29875123** |
| **SERPING1** | Reduced Expression Mutations | HAE-C1-INH | Serpin Family G Member 1 (SERPING1), enocdes for the C1-Inhibitor, C1-INH. C1INH is a highly glycosylated plasma protein that inhibits the C1 protein complex, FXIIa, and FXIIf, which in turn preventing excess complement and bradykinin production. Mutations that decresae expression of SERPING1 5-30% result is the leading cause of Hereditary Angioedema (HAE). Symptoms of HAE include Edema, GI symptoms, respiratory distress, predisposition to thrombotic events, painless non-itchy rash, tingling skin, skin tightness, fatigue, irritability sudden mood changes, anxiety. | **24552232, 30592059** |
| **F12** | Overexpression Mutation | HAE-FXII | Coagulation Factor XII (F12), enocdes for the complement activator factor FXII. FXII is a serum glycoprotein that initiates blood coagulation, fibrinolysis, and the generation of bradykinin. Mutations that increase expression of F12 are the second leading cause of HAE. | **33508266, 30592059** |
| **ANGPT1** | Reduced Expression Mutation | HAE | Angiopoietin 1 (ANGPT1), a key inhibitor of angiogenesis, for which loss of function germline mutations are an uncommon cause for HAE | **33508266** |
| **PLG** | c.988A>G (p.Lys330Glu) Mutation | HAE | Plasminogen (PLG), encodes for plasminogen, the inactive precursor of the enzyme plasmin, which triggers the fiber-lysis inhibiting fiber deposition. Patients with the PLG mutation c.988A>G (p.Lys330Glu) develop vascular leakage pathophysiology and HAE. | **28795768, 30809376** |
| **HS3ST6** | c.430A>T (p.Thr144Ser) Polymorphism | HAE | Heparan Sulfate-Glucosamine 3-Sulfotransferase 6 (HS3ST6), encodes for a protein required for the last step heparan sulfate biosynthesis of heparan sulfate. Patients with the HS3ST6 c.430A>T (p.Thr144Ser) polymorphism have increased vasculogenesis, anticoagulant activation of antithrombin and develop HAE. | **33508266** |
| **KNG1** | Truncation Variant | HAE | The Kininogen 1 (KNG1) mRNA is cleaved into transcripts that either translate for the high molecular weight kininogen (HMWK) or the low molecular weight (LMWK). Mutation in KNG1 that favors HMWK cleavage, results in increased bradykinin-production resulting in excess complement mediated HAE | **31087670, 24552232** |
| **MYOF** | Frameshift mutation | HAE | Myoferlin (MYOF), encodes for myoferlin, which localizes to the VEGFR-2 protein to act as a growth factor for vascular endothelial cells to induce pathological angiogenesis with vessel proliferation and migration are associated with cardiomyopathy | **33508266** |

**Table S4. Characteristics of EPICC participants included in analyses.**

| | Acute COVID (N=221) | Control (N=99) | Total (N=320) | *P value*[a] |
|---|---|---|---|---|
| **Gender** | | | | < 0.01 |
| Female | 75 (33.9%) | 53 (53.5%) | 128 (40.0%) | |
| Male | 146 (66.1%) | 46 (46.5%) | 192 (60.0%) | |
| **Race** | | | | 0.32 |
| White | 104 (47.1%) | 52 (52.5%) | 156 (48.8%) | |
| Hispanic or Latino | 57 (25.8%) | 19 (19.2%) | 76 (23.8%) | |
| Black | 34 (15.4%) | 20 (20.2%) | 54 (16.9%) | |
| Others | 26 (11.8%) | 8 (8.1%) | 34 (10.6%) | |
| **Age group** | | | | < 0.01 |
| <18 | 0 (0.0%) | 3 (3.0%) | 3 (0.9%) | |
| 18-44 | 98 (44.3%) | 56 (56.6%) | 154 (48.1%) | |
| 45-64 | 83 (37.6%) | 31 (31.3%) | 114 (35.6%) | |
| 65+ | 40 (18.1%) | 9 (9.1%) | 49 (15.3%) | |
| **Charlson Comorbidity Index** | | | | 0.24 |
| 0 | 109 (49.3%) | 54 (54.5%) | 163 (50.9%) | |
| 1-2 | 58 (26.2%) | 30 (30.3%) | 88 (27.5%) | |
| 3-4 | 29 (13.1%) | 6 (6.1%) | 35 (10.9%) | |
| >5 | 25 (11.3%) | 9 (9.1%) | 34 (10.6%) | |
| **Severity** | | | | < 0.01 |
| Hospitalized | 100 (45.2%) | 20 (20.2%) | 120 (37.5%) | |
| Outpatient | 121 (54.8%) | 79 (79.8%) | 200 (62.5%) | |
| **Variants** | | | | |
| Pre Delta | 186 (84.2%) | 0 | 186 (84.2%) | |
| Delta period[b] | 35 (15.8%) | 0 | 35 (15.8%) | |

[a]*n x k Fisher's Exact Test*

[b]*July 1st 2021 to December 31st 2021*

**Table S5: Genotype of EPICC participants included in analyses.**

| | Acute COVID (N=221) | Control (N=99) | Total (N=320) |
|---|---|---|---|
| **PANGOLIN** | | | |
| B.1.2 | 16 (11.3%) | 0 | 16 (11.3%) |
| B.1 | 12 (8.5%) | 0 | 12 (8.5%) |
| B.1.243 | 7 (5.0%) | 0 | 7 (5.0%) |
| B.1.369 | 7 (5.0%) | 0 | 7 (5.0%) |
| B.1.1.7 | 6 (4.3%) | 0 | 6 (4.3%) |
| AY.100 | 4 (2.8%) | 0 | 4 (2.8%) |
| AY.103 | 3 (2.1%) | 0 | 3 (2.1%) |
| AY.25 | 3 (2.1%) | 0 | 3 (2.1%) |
| B.1.1.207 | 3 (2.1%) | 0 | 3 (2.1%) |
| B.1.617.2 | 3 (2.1%) | 0 | 3 (2.1%) |
| B.1.429 | 3 (2.1%) | 0 | 3 (2.1%) |
| B.1.637 | 2 (1.4%) | 0 | 2 (1.4%) |
| B.1.1.519 | 2 (1.4%) | 0 | 2 (1.4%) |
| B.1.1.222 | 2 (1.4%) | 0 | 2 (1.4%) |
| AY.3 | 2 (1.4%) | 0 | 2 (1.4%) |
| AY.119 | 2 (1.4%) | 0 | 2 (1.4%) |
| B.1.1 | 2 (1.4%) | 0 | 2 (1.4%) |
| B.1.206 | 2 (1.4%) | 0 | 2 (1.4%) |
| B.1.234 | 2 (1.4%) | 0 | 2 (1.4%) |
| B.1.351 | 2 (1.4%) | 0 | 2 (1.4%) |
| B.1.355 | 2 (1.4%) | 0 | 2 (1.4%) |
| B.1.355,B.1.369 | 2 (1.4%) | 0 | 2 (1.4%) |
| A.27 | 1 (0.7%) | 0 | 1 (0.7%) |
| A.3 | 1 (0.7%) | 0 | 1 (0.7%) |
| AY.103,AY.116 | 1 (0.7%) | 0 | 1 (0.7%) |
| AY.103,AY.44 | 1 (0.7%) | 0 | 1 (0.7%) |
| AY.103,B.1 | 1 (0.7%) | 0 | 1 (0.7%) |
| AY.103,B.1.617.2 | 1 (0.7%) | 0 | 1 (0.7%) |
| AY.110 | 1 (0.7%) | 0 | 1 (0.7%) |
| AY.116 | 1 (0.7%) | 0 | 1 (0.7%) |
| AY.117 | 1 (0.7%) | 0 | 1 (0.7%) |
| AY.118 | 1 (0.7%) | 0 | 1 (0.7%) |
| AY.118,B.1.617.2 | 1 (0.7%) | 0 | 1 (0.7%) |
| AY.14,B.1.617.2 | 1 (0.7%) | 0 | 1 (0.7%) |
| AY.25,B.1.617.2 | 1 (0.7%) | 0 | 1 (0.7%) |
| AY.25.1 | 1 (0.7%) | 0 | 1 (0.7%) |
| AY.25.1,B.1.617.2 | 1 (0.7%) | 0 | 1 (0.7%) |
| AY.44 | 1 (0.7%) | 0 | 1 (0.7%) |
| B.1,B.1.147,B.1.369 | 1 (0.7%) | 0 | 1 (0.7%) |
| B.1,B.1.162 | 1 (0.7%) | 0 | 1 (0.7%) |
| B.1,B.1.332 | 1 (0.7%) | 0 | 1 (0.7%) |

| | Acute COVID (N=221) | Control (N=99) | Total (N=320) |
|---|---|---|---|
| B.1,B.1.336 | 1 (0.7%) | 0 | 1 (0.7%) |
| B.1,B.1.428 | 1 (0.7%) | 0 | 1 (0.7%) |
| B.1.1.148 | 1 (0.7%) | 0 | 1 (0.7%) |
| B.1.1.291 | 1 (0.7%) | 0 | 1 (0.7%) |
| B.1.1.304 | 1 (0.7%) | 0 | 1 (0.7%) |
| B.1.1.318 | 1 (0.7%) | 0 | 1 (0.7%) |
| B.1.1.339 | 1 (0.7%) | 0 | 1 (0.7%) |
| B.1.1.380 | 1 (0.7%) | 0 | 1 (0.7%) |
| B.1.1.432 | 1 (0.7%) | 0 | 1 (0.7%) |
| B.1.1.7,B.1.2 | 1 (0.7%) | 0 | 1 (0.7%) |
| B.1.110.3 | 1 (0.7%) | 0 | 1 (0.7%) |
| B.1.110.3,B.1.369 | 1 (0.7%) | 0 | 1 (0.7%) |
| B.1.126 | 1 (0.7%) | 0 | 1 (0.7%) |
| B.1.139,B.1.2 | 1 (0.7%) | 0 | 1 (0.7%) |
| B.1.2,B.1.333 | 1 (0.7%) | 0 | 1 (0.7%) |
| B.1.2,B.1.369 | 1 (0.7%) | 0 | 1 (0.7%) |
| B.1.233 | 1 (0.7%) | 0 | 1 (0.7%) |
| B.1.236 | 1 (0.7%) | 0 | 1 (0.7%) |
| B.1.240 | 1 (0.7%) | 0 | 1 (0.7%) |
| B.1.292 | 1 (0.7%) | 0 | 1 (0.7%) |
| B.1.298 | 1 (0.7%) | 0 | 1 (0.7%) |
| B.1.367 | 1 (0.7%) | 0 | 1 (0.7%) |
| B.1.400 | 1 (0.7%) | 0 | 1 (0.7%) |
| B.1.427,B.1.429 | 1 (0.7%) | 0 | 1 (0.7%) |
| B.1.427,B.1.637 | 1 (0.7%) | 0 | 1 (0.7%) |
| B.1.480 | 1 (0.7%) | 0 | 1 (0.7%) |
| B.1.556 | 1 (0.7%) | 0 | 1 (0.7%) |
| B.1.561 | 1 (0.7%) | 0 | 1 (0.7%) |
| B.1.565 | 1 (0.7%) | 0 | 1 (0.7%) |
| B.1.595 | 1 (0.7%) | 0 | 1 (0.7%) |
| B.1.605 | 1 (0.7%) | 0 | 1 (0.7%) |
| P.1.13 | 1 (0.7%) | 0 | 1 (0.7%) |
| Q.4 | 1 (0.7%) | 0 | 1 (0.7%) |
| NA | 80 (36.2%) | 99 (100%) | 179 (55.9%) |
